# Regulation of Interorganellar Ca^2+^ Transfer and NFAT Activation by the Mitochondrial Ca^2+^ Uniporter

**DOI:** 10.1101/2021.04.07.438854

**Authors:** Ryan E. Yoast, Scott M. Emrich, Xuexin Zhang, Ping Xin, Vikas Arige, Trayambak Pathak, J. Cory Benson, Martin T. Johnson, Natalia Lakomski, Nadine Hempel, Jung Min Han, Geneviève Dupont, David I. Yule, James Sneyd, Mohamed Trebak

**Affiliations:** Department of Cellular and Molecular Physiology, The Pennsylvania State University College of Medicine, 500 University Drive, Hershey, PA 17033, USA; Department of Pharmacology, The Pennsylvania State University College of Medicine, 500 University Drive, Hershey, PA 17033, USA; Department of Pharmacology and Physiology, University of Rochester, 601 Elmwood Ave, Rochester, NY 14642 USA; Laboratory of Biological Modeling, National Institute of Diabetes and Digestive and Kidney Diseases, National Institutes of Health, Bethesda, MD 20892, USA; Unité de Chronobiologie Théorique, Université Libre de Bruxelles, CP231, Boulevard du Triomphe, 1050, Brussels, Belgium; Department of Mathematics, The University of Auckland, 38 Princes Street, Auckland, 1010, New Zealand

## Abstract

Mitochondrial Ca^2+^ uptake is crucial for coupling receptor stimulation to cellular bioenergetics. Further, Ca^2+^ uptake by respiring mitochondria prevents Ca^2+^-dependent inactivation (CDI) of store-operated Ca^2+^ release-activated Ca^2+^ (CRAC) channels and inhibits Ca^2+^ extrusion to sustain cytosolic Ca^2+^ signaling. However, how Ca^2+^ uptake by the mitochondrial Ca^2+^ uniporter (MCU) shapes receptor-evoked interorganellar Ca^2+^ signaling is unknown. Here, we generated several cell lines with MCU-knockout (MCU-KO) as well as tissue-specific MCU-knockdown mice. We show that mitochondrial depolarization, but not MCU-KO, inhibits store-operated Ca^2+^ entry (SOCE). Paradoxically, despite enhancing Ca^2+^ extrusion and promoting CRAC channel CDI, MCU-KO increased cytosolic Ca^2+^ in response to store depletion. Further, physiological agonist stimulation in MCU-KO cells led to enhanced frequency of cytosolic Ca^2+^ oscillations, endoplasmic reticulum Ca^2+^ refilling, NFAT nuclear translocation and proliferation. However, MCU-KO did not affect inositol-1,4,5-trisphosphate receptor activity. Mathematical modeling supports that MCU-KO enhances cytosolic Ca^2+^, despite limiting CRAC channel activity.

## Introduction

In addition to their well-established role in cellular energy production and metabolism, mitochondria play a critical role in cellular signaling pathways that regulate gene transcription, and cell survival and function (*1, 2*). In particular, mitochondria are active participants in cellular Ca^2+^ signaling(*3–8*). Mitochondrial Ca^2+^ uptake, which is driven by the steep voltage gradient across the inner mitochondrial membrane occurs through a protein complex containing the pore-forming mitochondrial Ca^2+^ uniporter (MCU) protein(*9, 10*). MCU forms a Ca^2+^-selective tetrameric channel in the inner mitochondrial membrane that is regulated by the gate-keeping function of Ca^2+^-binding MICU1/2 protein dimers(*11–14*). MICU1/2 dimers keep the MCU channel closed under resting levels of free cytosolic Ca^2+^. Increased Ca^2+^ concentration in the vicinity of MCU and Ca^2+^ binding to the EF-hand domains of MICU1/2 disinhibits MCU channels and enhances mitochondrial Ca^2+^ uptake(*13, 14*). Ca^2+^ extrusion from the mitochondrial matrix to the cytosol occurs through independent transporters, which include the mitochondrial Na^+^/Ca^2+^ exchanger NCLX(*15*) and possibly the Ca^2+^/H^+^ exchanger Letm1(*16*). The gatekeeping of MCU channel activity by MICU1/2 is relieved only when cytosolic Ca^2+^ concentration is high (above ∼1.3 µM and above 500 nM when only MICU1 is present(*13*), but see also (*14*) where calculated Kd for Ca^2+^ binding to MICU1/2 dimers is ∼650 nM). As such, mitochondrial Ca^2+^ uptake is thought to take place at specialized microdomains where cytosolic Ca^2+^ concentrations are high. Such microdomains include the mitochondria-associated membranes (MAMs) where the ER membrane is within 10-30 nm from the outer mitochondrial membrane and Ca^2+^ is transferred to mitochondria through closely apposed inositol-1,4,5-trisphosphate receptors (IP_3_R) channels within ER membranes(*17*).

Activation of plasma membrane (PM) receptors that couple to isoforms of phospholipase C (PLC) by hormones, neurotransmitters and growth factors causes the breakdown of membrane-associated phosphatidylinositol-4,5-bisphosphate (PIP_2_) into two second messengers: the membrane-bound diacylglycerol (DAG) and the diffusible IP_3_(*18*). Activation of IP_3_R channels located in the endoplasmic reticulum (ER) causes ER Ca^2+^ store depletion. Upon Ca^2+^ store depletion, ER-resident stromal interaction molecules 1 and 2 (STIM1/2) undergo a conformational change and move to ER-PM junctional spaces where they form puncta and activate Orai channels mediating the highly Ca^2+^ selective, store-operated, Ca^2+^ release-activated Ca^2+^ (CRAC) current(*19–21*). Under conditions of stimulation with low concentrations of receptor agonists believed to represent physiological levels of receptor stimulation, cytosolic Ca^2+^ signals manifest as regenerative Ca^2+^ oscillations, which result from cycles of Ca^2+^ release through IP_3_R and concomitant bursts of CRAC channel activities that, depending on the cell type, can either directly sustain Ca^2+^ oscillations or replenish the depleted ER stores to sustain IP_3_R-driven Ca^2+^ oscillations(*22–24*). Local Ca^2+^ entry through Orai/CRAC channels leads to the activation of isoforms of the nuclear factor for activated T-cells (NFAT) transcription factors(*25*), which activate gene programs that control various cell functions, including proliferation and metabolism(*26, 27*).

As Ca^2+^ entry through CRAC channels accumulates on the cytosolic side of the plasma membrane, it mediates inhibition of CRAC channels through several feedback mechanisms. First, there is fast Ca^2+^-dependent inactivation (CDI) that involves Ca^2+^ binding to inhibitory sites within few nm of the mouth of CRAC channels (*28–31*). Mitochondria locate within 100-500 nm from the plasma membrane (PM) and because of their size, they would be presumably excluded from the tight ER-PM junctional sites that span ∼10-25nm where STIM and Orai co-aggregate. Therefore, it is unlikely that mitochondria can affect fast CDI of CRAC channels in any meaningful way. Second, there is slow CDI that is triggered by store refilling(*32*), which is due to reversal of STIM puncta independently of mitochondria. Third, there is another form of slow CDI that occurs independently of store refilling (e.g. in the presence of thapsigargin) (*32*). This store-independent slow CDI is mediated by Ca^2+^-binding sites that are located ∼100 nm or more from CRAC channels (*32*), and was proposed to be mediated by Ca^2+^-calmodulin (CaM)-mediated dissociation of STIM1 puncta and STIM1/Orai1 complexes(*33*). This latter form of slow CDI could be accounted for by mitochondrial Ca^2+^ buffering at a distant microdomain where CaM is located. Previous studies pre-dating the discovery of MCU and STIM/Orai have used drugs such as CCCP and antimycin A1, which inhibit mitochondrial Ca^2+^ uptake by altering mitochondrial respiration or depolarizing its membrane, to propose that mitochondria can buffer Ca^2+^ to alleviate slow CDI and sustain SOCE in Jurkat and primary T-cells(*34, 35*). A more recent study utilized MCU knockdown with siRNA in rat basophilic leukemia (RBL) cells and suggested that mitochondrial Ca^2+^ buffering specifically by MCU is required for sustaining the activity of IP_3_R, CRAC channels and agonist-evoked Ca^2+^ oscillations(*36*).

Here, we have investigated the role of MCU in regulating cytosolic, ER and mitochondrial Ca^2+^ dynamics using an arsenal of cell lines and primary cells from different species and tissues. We utilized CRISPR/Cas9 to produce several clones of MCU knockout in cultured cells and used the Cre-LoxP system to generate populations of primary cells from tissue-specific MCU knockdown mice. We show that despite promoting slow CDI of CRAC channels and enhancing cytosolic Ca^2+^ extrusion, MCU deletion leads to enhanced cytosolic Ca^2+^ in response to passive store depletion. MCU-KO also enhanced the frequency of cytosolic Ca^2+^ oscillations in response to agonist stimulation without affecting IP_3_R-mediated Ca^2+^ puffs. MCU-KO causes enhanced ER Ca^2+^ refilling and increased nuclear translocation of NFAT isoforms, and increased lymphocyte proliferation. Mathematical modeling is consistent with Ca^2+^ buffering by MCU as an important regulator of interorganellar Ca^2+^ transfer that fine-tunes cytosolic Ca^2+^ signaling independently of IP_3_Rs and CRAC channels. Our data provide critical insights into the role of MCU and mitochondrial Ca^2+^ uptake in cellular signaling. Despite the critical function of MCU-mediated mitochondrial Ca^2+^ uptake in inhibiting CRAC channel slow CDI and limiting Ca^2+^ extrusion, MCU deletion leads to a paradoxical enhancement of cytosolic Ca^2+^ upon receptor stimulation. Thus, the inability of mitochondria to store its “allocated share” of Ca^2+^ in the absence of MCU largely offsets the effects on Ca^2+^ extrusion and CRAC channel activity. The net result of MCU-KO is enhanced ER Ca^2+^ refilling and increased cytosolic Ca^2+^ signaling.

## Results

### MCU knockout enhances cytosolic Ca^2+^ despite promoting Ca^2+^ extrusion and CRAC channel slow CDI

We performed CRISPR/cas9 gene knockout of MCU in six cell lines from different tissues and species (**Fig. 1**). A seventh cell line (Hela) and its corresponding MCU CRISPR/Cas9 knockout clone were kindly provided by Dr. Suresh Joseph (Thomas Jefferson University) (**Fig. S1**). The cell lines are: human HeLa, embryonic kidney (HEK293), colorectal cancer HCT116 and DLD1, and Jurkat T-cells; rat basophilic leukemia (RBL-1) mast cells; and mouse A20 B-cells. These MCU knockout cell lines provided a clean background to analyze the role of mitochondrial Ca^2+^ uptake in shaping intracellular Ca^2+^ signaling. To alleviate potential off-target effects of CRISPR/Cas9 KO, we studied two independent MCU-KO clones for each cell line (except for HeLa cells, of which we obtained only 1 clone). These clones were identified, validated for MCU-KO with genomic sequencing, Western blot and mitochondrial Ca^2+^ measurements, and assayed in parallel to their corresponding parental controls. Absence of the MCU protein in knockout cells was documented by Western blot (**Fig. 1A, E, I, C, G, K**). Functional knockout of MCU was documented by simultaneous Ca^2+^ and mitochondrial membrane potential fluorescence measurements in a permeabilized cell system (**Fig. S2**). The addition of a bolus 10 µM Ca^2+^ to the bath of permeabilized parental cells resulted in rapid mitochondrial uptake, which did not occur in MCU-KO cells, demonstrating that acute mitochondrial Ca^2+^ uptake was abrogated in MCU-KO cells (**Fig. S2**).

**Figure 1.**
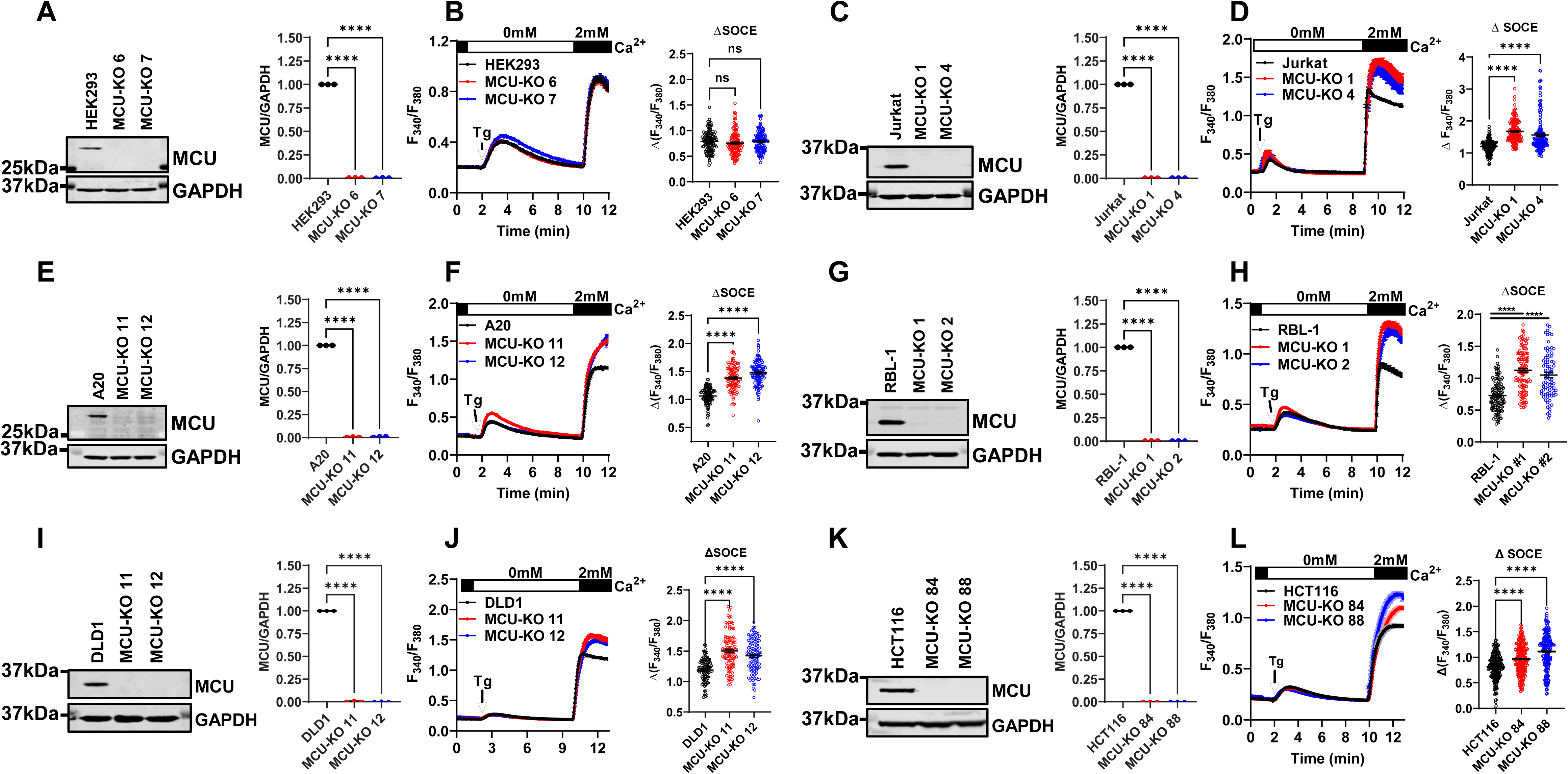
MCU-KO enhances cytosolic Ca^2+^ upon thapsigargin stimulation. Western blot documenting MCU protein knockout in two independent CRISPR/Cas9 MCU-KO clones of HEK293 cells and quantification of MCU protein band densitometry relative to GAPDH from three independent experiments (**A**). Ca^2+^ measurements in HEK293 parental cells and MCU-KO clones in response to stimulation with 2 µM thapsigargin (Tg) in Ca^2+^-free buffer followed by restoration of 2 mM extracellular Ca^2+^ to determine the magnitude of SOCE (**B**). Quantification of SOCE (Δ SOCE) from at least three independent experiments are also shown (**B**). Identical experiments to (**A, B**) were performed in human Jurkat T-cells (**C, D**), mouse A20 B-cells (**E, F**), rat RBL-1 mast cells (**G, H**), human DLD1 colorectal carcinoma cells (**I, J**) and human HCT116 colorectal carcinoma cells (**K, L**).

The predominant receptor-activated Ca^2+^ entry pathway in all seven non-excitable cell lines considered herein is the store-operated Ca^2+^ entry (SOCE) pathway, which biophysically manifests as the Ca^2+^ release-activated Ca^2+^ (CRAC) current encoded by STIM1/2 and Orai1/2/3 proteins. To maximally activate SOCE, we irreversibly blocked the Sarcoplasmic/ER Ca^2+^ ATPase (SERCA) using thapsigargin (Tg; 2 µM), which causes passive depletion of ER Ca^2+^ stores (recorded in Ca^2+^-free bath solutions) and activation of SOCE, which manifests upon addition of 2 mM Ca^2+^ to the extracellular bath (**Fig. 1B, F, J, D, H, L**). Surprisingly, the use of this protocol revealed that MCU-KO led to a significant increase in apparent SOCE in all cell lines, with the exception of MCU-KO HEK293 cells where SOCE was increased in some runs but not significantly different from that of the parental HEK293 cells when all cells were averaged (**Fig. 1B**; see also **Fig. S5A**). This notable difference between HEK293 cells and the other six cell lines will be addressed further below. Western blots showed no change in STIM1 protein levels upon MCU-KO in either HEK293 cells or Jurkat cells (**Fig. S3**). Quantitative PCR analysis revealed no significant compensatory up/downregulation of all five STIM/Orai proteins in HEK293 or Jurkat MCU-KO cells (**Fig. S4**).

Previous reports showed that abrogation of the ability of mitochondria to buffer cytosolic Ca^2+^, through dissipation of the mitochondrial membrane potential by the protonophore carbonyl cyanide *m*-chlorophenylhydrazone (CCCP), or blockade of complex III and complex V of the electron transport chain (ETC) with Antimycin A1 and oligomycin, led to inhibition of SOCE(*35*). CRAC currents were also inhibited by these drugs, when recorded with either the perforated patch-clamp technique or the whole-cell technique with pipette solutions containing a relatively low Ca^2+^ buffer (1.2 mM EGTA) but including metabolites that promote energized mitochondria (*34*). These findings showed that a healthy mitochondrial membrane potential supports mitochondrial Ca^2+^ uptake, which in turn limits slow Ca^2+^-dependent inhibition (CDI) of CRAC channels to sustain SOCE.

Therefore, we performed SOCE measurements in response to thapsigargin in parental HEK293, Jurkat and RBL-1 cells and their MCU-KO counterparts and tested the effect of 5 µM of the protonophore trifluoromethoxy carbonylcyanide phenylhydrazone (FCCP) on SOCE. As seen in **Fig. S5A-C**, when added after SOCE was initiated, FCCP significantly inhibited SOCE in both parental cells and their MCU-KO counterparts. The inhibition of SOCE in parental Jurkat (**Fig. S5B**) and RBL-1 (**Fig. S5C**) cells was slower than in their MCU-KO counterparts and was preceded by a transient increase in Ca^2+^, presumably reflecting bigger mitochondrial Ca^2+^ stores in the parental cells. In HEK293 cells, the rate of SOCE inhibition by FCCP was similar between wildtype and MCU-KO cells (**Fig. S5A**), suggesting a bigger contribution of mitochondria as a reservoir for Ca^2+^ in Jurkat and RBL-1 cells compared to HEK293 cells. Addition of 1 µM ionomycin to HEK293 and Jurkat cells (red arrow; **Fig. S5A, B**) to release any remaining Ca^2+^ from thapsigargin-independent stores (presumably mitochondria) showed a small mitochondrial Ca^2+^ release in wildtype Jurkat cells, but not in wildtype HEK293 cells. We subsequently added 10 µM ionomycin (blue arrow) in the presence of 2 mM Ca^2+^ to confirm that maximal Fura2 signal is equal between MCU-KO and parental cells. Overall, these data argue that inhibition of SOCE by FCCP cannot be explained simply by lack of mitochondrial Ca^2+^ buffering through MCU. One recurring observation in our recordings is the enhanced rate of decline of SOCE steady-state plateaus in Jurkat and RBL-1 MCU-KO cells compared to parental cells (**Fig. S5B, C**), suggesting that mitochondrial Ca^2+^ buffering by MCU inhibits PM Ca^2+^ ATPase (PMCA)-mediated Ca^2+^ extrusion, as shown previously in Jurkat T-cells (*37*). Indeed, Inhibition of SOCE with 5 µM Gd^3+^ revealed that the rate of Ca^2+^ extrusion is significantly faster in MCU-KO cells compared to Jurkat (**Fig. S6A, B**) and RBL-1 parental cells (**Fig. S6C, D**).

We then sought to determine whether increased SOCE upon MCU-KO is due to increased net CRAC currents across the plasma membrane by using the whole-cell patch clamp technique. We chose to record from two widely studied cell lines that produce the largest native CRAC currents, namely Jurkat and RBL-1 cells. We recorded both Ca^2+^ CRAC currents in 20mM Ca^2+^ bath solutions and performed rapid switches to divalent free (DVF) bath solutions to record Na^+^ CRAC currents, which are larger in size. We measured whole-cell CRAC currents from Jurkat and RBL-1 cells under two different conditions. The first condition uses a pipette solution with a low buffering capacity (1 mM EGTA) that includes 30 µM IP_3_ to empty the stores and contains a cocktail of mitochondria-energizing metabolites (see Methods) that might reveal mitochondrial effects on CRAC channel slow CDI. The second condition uses a pipette solution with a very strong buffer (50 mM BAPTA), that would negate any potential effect of mitochondrial Ca^2+^ buffering on CRAC currents. Native Ca^2+^ and Na^+^ CRAC currents under both weak and strong buffer conditions in MCU-KO clones of both Jurkat and RBL-1 cells were statistically of similar magnitude compared to currents from their corresponding parental cells (**Fig. S7**). To accurately determine if MCU deletion alters store-independent CRAC channel slow CDI, we used an established patch clamp protocol where Jurkat cells are pre-incubated with thapsigargin in the absence of extracellular Ca^2+^ and I_CRAC_ is recorded on introduction of 20 mM Ca^2+^ to the bath solution (*32*). I_CRAC_ was recorded in both Jurkat MCU-KO and parental cells with a pipette solution containing a low buffer (1.2 mM EGTA, 0.66 mM Ca^2+^ (*32*); see methods) either with or without supplementation with the mitochondria-energizing cocktail (**Fig. S8**). Interestingly, even in the absence of the mitochondria-energizing cocktail, MCU-KO cells showed enhanced slow CDI compared to control cells (**Fig. S8A-C**). Furthermore, when the pipette solution included the mitochondria-energizing cocktail, I_CRAC_ slow CDI of parental Jurkat cells was inhibited on average by ∼90%, while I_CRAC_ slow CDI in MCU-KO cells remained unchanged (**Fig. S8D-F**). CDI recordings with and without mitochondria-energizing cocktail in each individual wildtype Jurkat and MCU-KO cell are shown in **Fig. S9** and **Fig. S10**, respectively.

Taken together, these data indicate that although abrogation of MCU-mediated mitochondrial Ca^2+^ uptake promotes Ca^2+^ extrusion and CRAC channel CDI, its net effect is an increase, rather than a decrease, in cytosolic Ca^2+^ levels. Our findings suggest that preventing mitochondria from buffering its “allocated share” of cytosolic Ca^2+^ could lead to accelerated ER Ca^2+^ refilling, and enhanced frequency of agonist-evoked cytosolic Ca^2+^ oscillations, NFAT activation and cell proliferation. These concepts are explored further below.

### MCU knockout enhances the frequency of cytosolic Ca^2+^ oscillations without affecting IP_3_R activity

We determined the effect of MCU-KO on Ca^2+^ signaling triggered by physiological agonist stimulation. To induce cytosolic Ca^2+^ oscillations, Jurkat cells were stimulated with a relatively low concentration (0.125 µg/mL) of anti-CD3 antibody, which is commonly used to activate the T-cell receptor (TCR) and drive clonal expansion and interleukin 2 (IL-2) production. Under these conditions, essentially all Jurkat cells that responded displayed regenerative Ca^2+^ oscillations (**Fig. 2A, B**) with MCU-KO Jurkat cells displaying enhanced frequency of Ca^2+^ oscillations compared to the parental cells (**Fig. 2A-C**). Further, Ca^2+^ oscillations in MCU-KO Jurkat cells were of longer duration compared to those of the parental cells (**Fig. 2D, E**).

**Figure 2.**
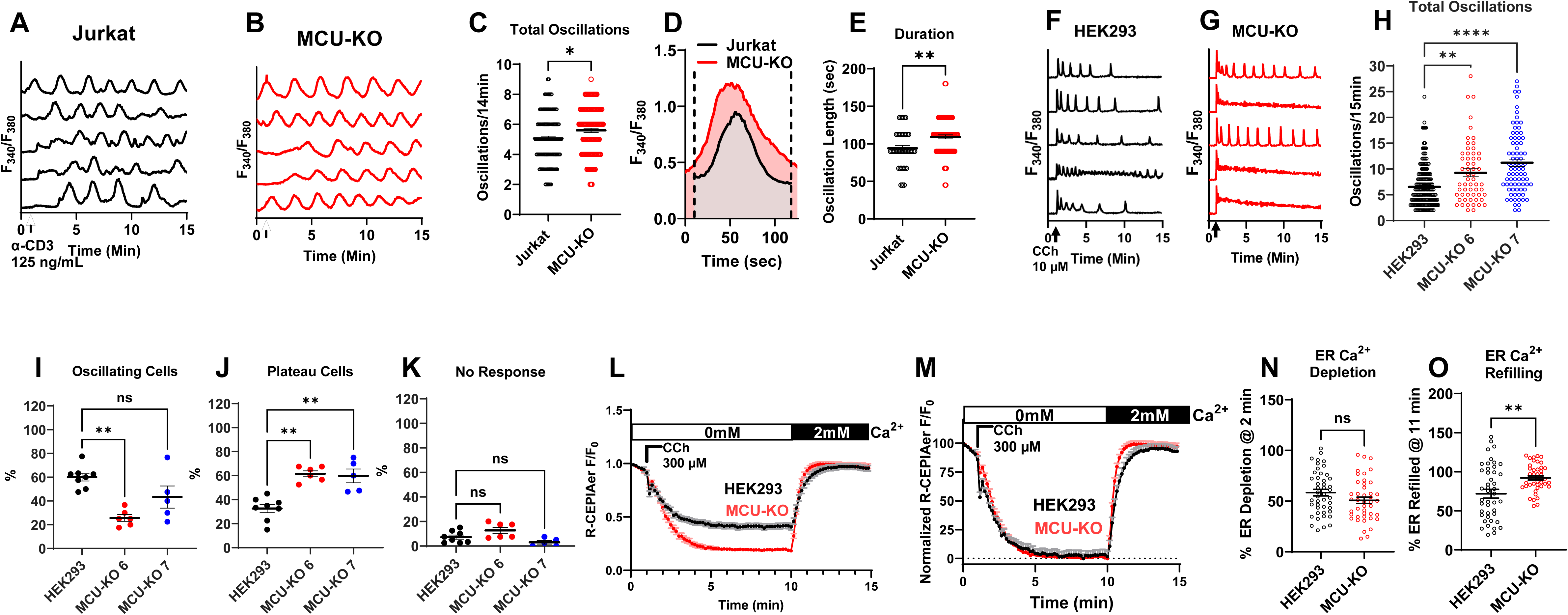
MCU-KO enhances the frequency of Ca^2+^ oscillations and ER refilling. (**A-E**) Ca^2+^ oscillations were elicited in wildtype Jurkat cells (**A**) and their MCU-KO counterparts (**B**) by stimulation with 0.125 µg/mL of anti-CD3 antibody in Ca^2+^-containing (2 mM) buffer. The number of Ca^2+^ oscillations/14 min (**C**), overlaid representative spikes (**D**) and quantification of oscillation duration in sec (**E**) in Jurkat cells and their MCU-KO variant are represented. (**F-K**) Ca^2+^ oscillations were elicited in wildtype HEK293 cells (**F**) and their MCU-KO counterparts (**G**) by stimulation with10 µM carbachol (CCh) in Ca^2+^-containing (2 mM) buffer. (**H**) Quantification of the number of oscillations/14 min in wildtype HEK293 cells and their MCU-KO clones #6 and #7. Quantification of the percentage of HEK293 cells and MCU-KO cells that respond with either repetitive Ca^2+^ oscillations (**I**), or sustained Ca^2+^ plateaus (**J**) as well as cells that show no response (**K**) from several independent runs. (**L-N**), Direct measurements of ER Ca^2+^ levels using CEPIAer in HEK293 cells and their MCU-KO counterparts. Cells were stimulated with 300 µM CCh in Ca^2+^-free buffer to elicit ER Ca^2+^ depletion followed by CCh washout and replenishment of 2mM Ca^2+^ to measure ER refilling (**L**). Data in (**L**) are normalized in (**M**) and rates of ER depletion (**N**) and refilling (**O**) were quantified from several cells and statistically analyzed.

Stimulation of HEK293 with a low concentration of carbachol (10 µM CCh), in the presence of 2 mM extracellular Ca^2+^, evoked regenerative Ca^2+^ oscillations in the majority of cells (**Fig. 2F, I-K**). Note that HEK293 cells was the only cell line studied where SOCE in response to thapsigargin was not significantly altered by MCU-KO. Yet, MCU-KO HEK293 cells showed a higher proportion of cells responding with a Ca^2+^ plateau at the expense of cells responding with repetitive Ca^2+^ oscillations compared to their parental counterparts (**Fig. 2I-K**). Further, when oscillating cells are considered MCU-KO HEK293 cells showed a higher frequency of cytosolic Ca^2+^ oscillations than their parental counterparts (**Fig. 2H**), suggesting that MCU-KO leads to higher cytosolic Ca^2+^. It is not obvious why the cytosolic Ca^2+^ signal in response to thapsigargin is similar between MCU-KO HEK293 cells and their parental counterparts (despite clear differences in Ca^2+^ oscillations triggered by agonist). We reasoned that differences in mitochondrial and/or cytosolic volumes between these different cells might play a role. We performed high resolution three-dimensional multispectral confocal microscopy to generate volumetric reconstructions of the cytosol and mitochondria of Jurkat, A20, RBL-1 and HEK293 cell lines (**Fig. S11A-E**). Our volumetric measurements showed that HEK293 cells have a higher cytosolic and cell volume compared to all other cell lines (**Fig. S11A-C**). However, we found no obvious correlation between the mitochondrial volume or the mitochondria/cell ratio and the differences in SOCE in response to thapsigargin between parental and MCU-KO cells (**Fig. S11D, E**). Interestingly, side by side comparisons of mitochondrial Ca^2+^ extrusion between wildtype HEK293, Jurkat, A20 and RBL-1 cells using the permeabilized cell preparation showed that HEK293 cells have significantly enhanced mitochondrial Ca^2+^ extrusion compared to all other cell types (**Fig. S11F-I**). The enhanced mitochondrial Ca^2+^ extrusion in HEK293 cells suggests a moderate role of mitochondria in buffering cytosolic Ca^2+^ in these cells (see also **Fig. S5A** where FCCP effect is similar between parental and MCU-KO HEK293 cells). This might contribute to the blunting of small differences in the Ca^2+^ signal between MCU-KO and parental HEK293 cells under the thapsigargin protocol. Additional studies are needed to resolve this issue.

We also performed measurements of ER Ca^2+^ stores in MCU-KO and wildtype HEK293 cells by transiently expressing the genetically encoded Ca^2+^ sensor R-CEPIAer. Treatment of cells with supramaximal concentration of carbachol (300 µM CCh) in Ca^2+^-free solution caused a significantly bigger drop of CEPIAer fluorescence in MCU-KO cells compared to parental cells (**Fig. 2L**), suggesting a higher ER Ca^2+^ content in MCU-KO cells compared to parental cells. These CEPIAer fluorescence measurements are represented as normalized data in **Fig. 2M**. We did not resolve a difference in the rate of ER Ca^2+^ depletion between wildtype and MCU-KO HEK293 cells (**Fig. 2N**). However, the rate of ER Ca^2+^ refilling after CCh wash-off and addition of 2mM Ca^2+^ to the bath was significantly faster in MCU-KO HEK293 cells (**Fig. 2O**), suggesting that MCU function reduces cytosolic Ca^2+^ and the subsequent efficiency of ER Ca^2+^ refilling.

To determine whether enhanced Ca^2+^ oscillatory frequency of MCU-KO cells is due to increased IP_3_R activity, we used TIRF microscopy to measure the elementary Ca^2+^ signals called Ca^2+^ puffs, which are mediated by IP_3_Rs. We loaded parental HEK293 and MCU-KO cells with a Ca^2+^ dye and caged IP_3_ (ci-IP_3_/PM; see methods) and Ca^2+^ puffs were recorded at a rate of 166 frames/sec upon photolysis of ci-IP_3_. We detected Ca^2+^ puffs in both parental HEK293 and MCU-KO cells following photolysis of ci-IP_3_ (**Fig. S12**). However, the number of puffs and the number of puff sites recorded in either Ca^2+^-free or Ca^2+^-containing bath solutions were not significantly different between MCU-KO cells and parental HEK293 cells (**Fig. S12A, B, F, G**). The amplitude distribution of puffs did not differ between HEK293 and MCU-KO cells with most puffs ranging from 0.2 to 0.7 peak amplitude (**Fig. S12C, H**). The mean rise (r) and fall (f) time of the Ca^2+^ puffs were also similar between HEK293 and MCU-KO cells (**Fig. S12D, I**), indicating that the fundamental biophysical properties of IP_3_R clusters were not different between these two groups of cells. Further, there was no difference between both groups of cells in the proportion of cells that showed a globalized Ca^2+^ signal within 60 sec (**Fig. S12E, J**). These data suggest that MCU knockout does not alter IP_3_R channel activity.

### MCU knockout enhances NFAT1/4 activation

Given IP_3_R activity is unaltered by MCU-KO, the simplest explanation for enhanced Ca^2+^ oscillations in MCU-KO cells is that MCU-KO causes extra-mitochondrial Ca^2+^ build-up as mitochondria are unable to buffer their “allocated share” of cytosolic Ca^2+^. Hence, cytosol-ER-mitochondrial Ca^2+^ transfer would be backed up in MCU-KO cells, leading to faster ER refilling and/or increased net cytosolic Ca^2+^. We tested whether the net increase in global cytosolic Ca^2+^ we observed in MCU-KO cells is associated with enhanced NFAT activity, which is controlled by Ca^2+^ microdomains near CRAC channels. NFAT activity was significantly enhanced in MCU-KO cells of both HEK293 (**Fig. 3A-C**) and Jurkat cells (**Fig. 3D-F**) compared to their respective controls. We determined the effect of MCU-KO on NFAT4 nuclear translocation in HEK293 cells using a reporter NFAT4-GFP construct coupled to fluorescence microscopy (**Fig. 3A-C**). In Jurkat cells, we determined the effect of MCU-KO on *native* NFAT1 nuclear translocation using an NFAT1 specific antibody and ImageStream (**Fig. 3D-F**). Stimulation with 10 µM CCh caused a significantly bigger nuclear translocation of NFAT4 in MCU-KO HEK293 cells when compared to parental controls (**Fig. 3A-C**). Using a multispectral imaging flow cytometer (ImageStream) we tracked native NFAT1 nuclear translocation in response to ER store depletion with 2 µM thapsigargin for 30 min in populations of tens of thousands of MCU-KO Jurkat cells and their parental counterparts. MCU-KO Jurkat cells showed a significantly more robust nuclear translocation of endogenous NFAT1 compared to the parental Jurkat cells (**Fig. 3D-F**). These data suggest that the global cytosolic Ca^2+^ increase triggered by deletion of MCU leads to sufficient Ca^2+^ buildup at the vicinity of CRAC channels to enhance NFAT nuclear translocation.

**Figure 3.**
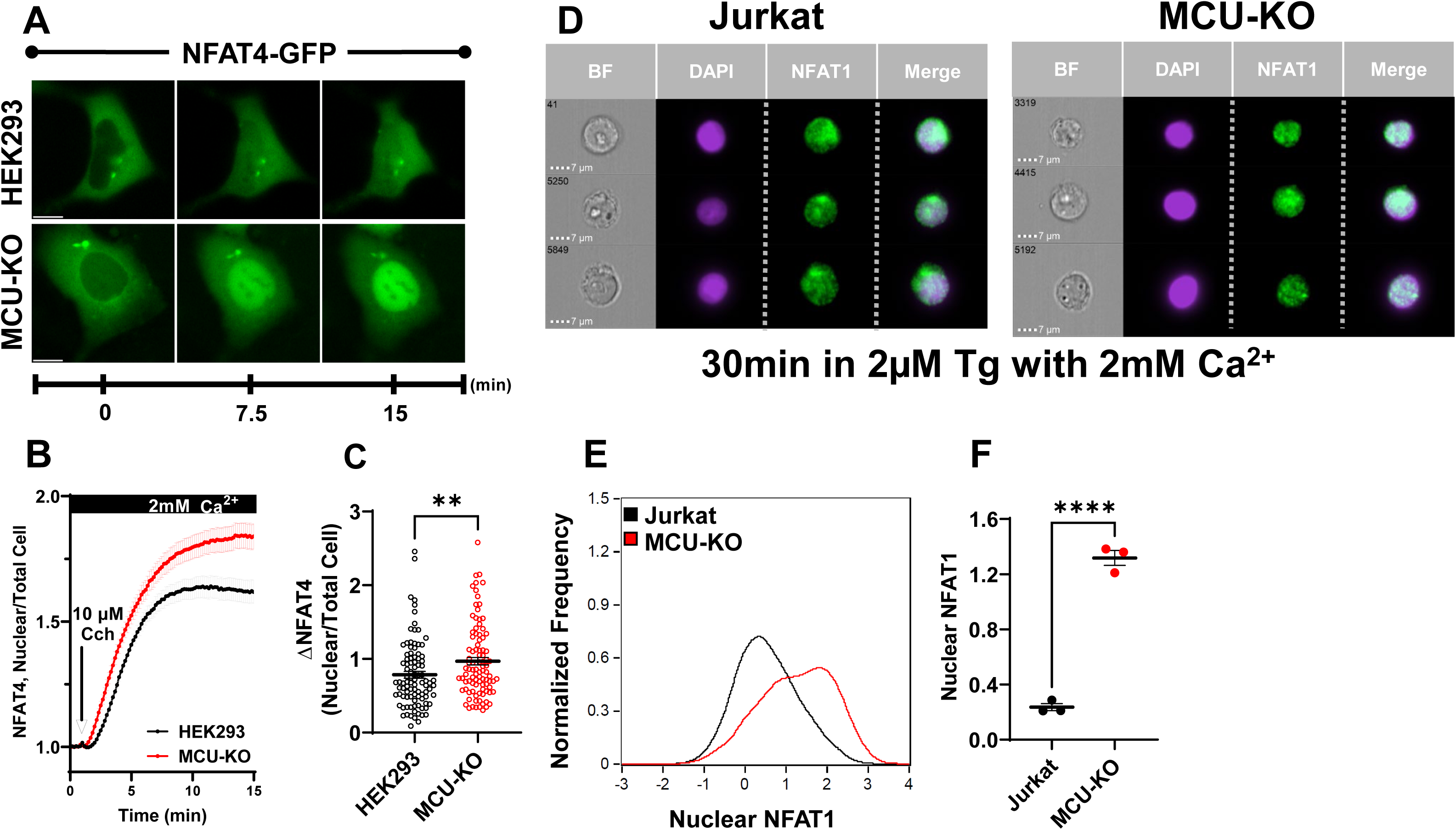
MCU-KO enhances NFAT nuclear translocation. (**A-C**) HEK293 cells and their MCU-KO counterparts were transfected with an NFAT4-GFP construct, stimulated with 10 µM carbachol (CCh) and nuclear translocation of NFAT4-GFP was monitored over time using a fluorescence microscope (**A**; see methods). The ratio of nuclear/total fluorescence of NFAT4-GFP (**B**) and difference between maximal and basal ratios (Δ NFAT4; **C**) are represented for both groups of cells and statistically analyzed. (**D-F**), ImageStream analysis of *native* NFAT1 nuclear translocation using flow cytometry in wildtype and MCU-KO Jurkat cells stimulated with 2 µM thapsigargin (Tg) for 30 min. Native NFAT1 was labeled with a fluorescently-tagged specific antibody and NFAT1 nuclear translocation was determined by co-localization of NFAT1 with DAPI nuclear staining (**D**). Histograms showing the distribution (**E**) and intensity (**F**) of nuclear NFAT1 fluorescence from three independent experiments with 10,000 cells/run in wildtype Jurkat and MCU-KO cells.

### Tissue-specific MCU knockdown in mice enhances lymphocyte SOCE and proliferation

To determine if the increase in cytosolic Ca^2+^ upon MCU-KO also occurs in primary cells and is not restricted to cell lines, we generated tissue specific knockdown (KD) mice for CD4^+^ T-cell and B-cells (see **Fig. S13** for mice genotyping). We used a 4-hydroxytamoxifen (4-OHT) inducible CD4 Cre line to generate mice with CD4^+^ T-cell specific MCU-KD (**Fig. 4A-G**). The B-cell specific MCU-KD mice were generated using the non-inducible MB1 Cre line (**Fig. 4H-M**). The LoxP-cre system led to reduction in MCU protein expression by ∼60-80% (**Fig. 4B-D, H, I**).

**Figure 4.**
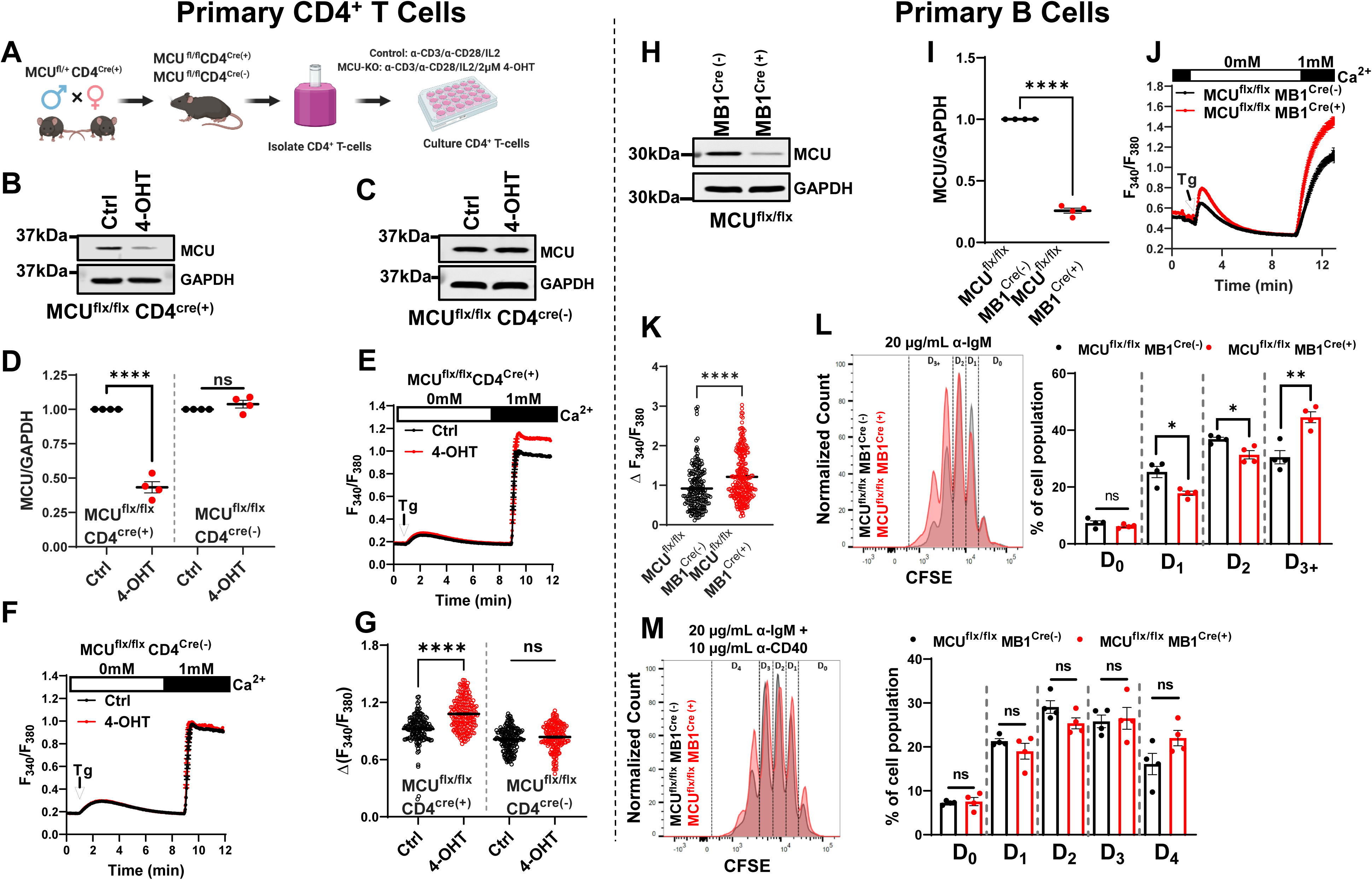
Tissue-specific MCU-knockdown (MCU-KD) in mice CD4^+^ T- and B-cells enhances SOCE and proliferation. (**A-G**) Primary CD4^+^ T-cells were isolated by negative selection (**A**) from spleens of MCU^flx/flx^ CD4^Cre(+)^ and MCU^flx/flx^ CD4^Cre(-)^ mice identified by genotyping. CD4^+^ T-cell specific MCU-KD was induced by treating the isolated cells *in vitro* with 4-OHT (**B-D**). After 4-OHT treatment, only CD4^+^ T cells from MCU^flx/flx^ CD4^Cre(+)^ mice show a reduction in MCU protein expression (**B, C**), which was quantified from four independent mice/group and statistically analyzed (**D**). (**F, H**), Fura2 Ca^2+^ measurements using the SOCE protocol in CD4^+^ T-cells isolated from MCU^flx/flx^ CD4^Cre(+)^ (**E**) and MCU^flx/flx^ CD4^Cre(-)^ (**F**) mice with and without 4-OHT treatment and maximal SOCE for each group of cells was statistically analyzed (**G**). (**H-M**), Primary B-cells were isolated by negative selection from spleens of MCU^flx/flx^ MB1^Cre(+)^ and MCU^flx/flx^ MB1^Cre(-)^ mice identified by genotyping. The MB1^Cre(+)^ line is not inducible and specific MCU-KD was documented on acutely isolated B-cells by Westerns (**H**) and MCU densitometry from four independent isolations was statistically analyzed (**I**). Fura2 Ca^2+^ measurements using the SOCE protocol in B-cells isolated from MCU^flx/flx^ MB1^Cre(+)^ and MCU^flx/flx^ MB1^Cre(-)^ mice (**J**) and maximal SOCE for each group of cells was statistically analyzed (**K**). (**L, M**), B-cell proliferation was determined by monitoring the dilution of the CFSE dye with each cell division (D0-D4) using flow cytometry after stimulation with either anti-IgM (**L**) or with anti-IgM + anti-CD40 (**M**).

Primary CD4^+^ T-cells were isolated with negative selection from spleens of MCU^flx/flx^ mice (MCU^flx/flx^ CD4^Cre(-)^) and MCU^flx/flx^ CD4^CreERT2^ mice (MCU^flx/flx^ CD4^Cre(+)^), activated *in vitro* using anti-CD3/anti-CD28 conjugated beads and mouse Interleukin-2 (IL-2), and treated with (Z)-4-Hydroxytamoxifen (4-OHT) to induce Cre activity for 96 hrs before experimentation. Control cells for both MCU^flx/flx^ CD4^Cre(+)^ and MCU^flx/flx^ CD4^Cre(-)^ conditions originated from the same mice but were not subjected to 4-OHT treatment. We verified that the administration of 4-OHT does not alter MCU protein levels in MCU^flx/flx^ CD4^Cre(-)^ mice (**Fig. 4C**). Stimulation of cells with 2 µM thapsigargin showed that only MCU-KD CD4^+^ T-cells, namely CD4^+^ T-cells isolated from MCU^flx/flx^ CD4^Cre(+)^ mice and treated with 4-OHT displayed significantly higher SOCE (**Fig. 4E-G**). Because the B-cell specific MB1Cre line is non-inducible, primary B-cells were isolated with negative selection from spleens of MCU^flx/flx^ mice either with MB1Cre (MCU^flx/flx^ MB1^Cre(+)^) or without (MCU^flx/flx^ MB1^Cre(-)^) and immediately used for Ca^2+^ imaging to measure SOCE. Stimulation of B-cells with 2 µM thapsigargin showed that B-cells isolated from MCU^flx/flx^ MB1^Cre(+)^ showed significantly higher SOCE compared to B-cells isolated from MCU^flx/flx^ MB1^Cre(-)^ mice (**Fig. 4J, K**).

The induction of MCU-KO with 4-OHT for five days in CD4^+^ T-cells from MCU^flx/flx^ CD4^Cre(+)^ mice precluded us from performing proliferation assays on these primary CD4^+^ T-cells. We did however perform proliferation assays on B-cells isolated from MCU^flx/flx^ MB1^Cre(+)^ and MCU^flx/flx^ MB1^Cre(-)^ mice. B-cell proliferation induced by weak BCR stimulation can be enhanced by Ca^2+^-independent signals such as co-stimulation with anti-CD40 or stimulation of toll-like receptors by lipopolysaccharides (LPS) (*38, 39*). Therefore, B-cells were stimulated either *via* the BCR with anti-IgM (**Fig. 4L**), co-stimulated with anti-IgM + anti-CD40 (**Fig. 4M**), LPS alone (**Fig. S14C**) or anti-CD40 alone as a control (**Fig. S12D**) using the CFSE dye (**Fig. S14A**). There were no differences in cell viability between different groups of cells and stimulatory conditions (**Fig. S14B**). Primary B-cells from MCU^flx/flx^ MB1^Cre(+)^ mice (MCU-KD B-cells) showed a significantly enhanced proliferation in response to BCR stimulation with anti-IgM compared to control MCU^flx/flx^ MB1^Cre(-)^ B-cells as documented by dilution of the CFSE dye with increased proportion of B-cells that underwent a third cycle of division (**Fig. 4L**). As expected, this difference in B-cell proliferation was not observed when cells were co-stimulated with anti-IgM + anti-CD40 (**Fig. 4M**) or when stimulated with LPS (**Fig. S14C**). Stimulation with anti-CD40 alone had marginal to no effect on B-cell proliferation (**Fig. S14D**).

### Mathematical modeling supports MCU-mediated regulation of cytosolic Ca^2+^

First, we tested how mitochondrial Ca^2+^ transport alone (i.e., without assuming the existence of microdomains that affect CRAC channel activity through CDI) can change the properties of cytosolic Ca^2+^ rise activated by passive store depletion with thapsigargin as we all as Ca^2+^ oscillations activated by agonist stimulation. We took an existing model of Ca^2+^ oscillations that has been validated against experimental data in different cell types(*23, 40, 41*), slightly simplified the description of the Ca^2+^ influx pathways, and added to it six different models, of increasing complexity, of mitochondrial Ca^2+^ transport. Model 1 assumes that the mitochondria act like a Ca^2+^ buffer, with highly simplified Ca^2+^ uptake by the MCU and Ca^2+^ extrusion *via* NCLX. Model 2 assumes that the MCU and NCLX fluxes are modelled in a more complex manner, following (*42*) (itself based on the earlier work of (*43, 44*)) but omitting the mitochondrial membrane potential and metabolism. Model 3 couples the model of (*42*), including the mitochondrial membrane potential and metabolism, to the Ca^2+^ oscillation model of(*23*), to obtain a system of 8 differential equations. Model 4 is the model of (*42*) with no changes to the equations or parameters. Model 5 extends model 3 to include MAMs, following (*45*). Model 6 is the model of (*45*) with no changes to parameters or equations.

When we simulate the effects of MCU-KO, NCLX-KO and MCU-KO + NCLX-KO on thapsigargin-activated cytosolic Ca^2+^ rise, we found that MCU-KO and MCU-KO + NCLX-KO cause similar increase in SOCE, while NCLX-KO leads to a slight decrease in SOCE (**Fig. 5A-E**), in agreement with our experimental data with MCU and previous work with NCLX(*46*). These simulations were performed for Model 2, 3 and 5 with three variations on Model 5 representing different percentages of total IP_3_Rs located within MAMs (10%, 50% or 70%; **Fig. 5C-E**). These thapsigargin simulations could not be performed for Models 1, 4 and 6. Model 1 models mitochondria as a simple Ca^2+^ buffer, which does not allow for properly simulating MCU/NCLX-KO experiments. Model 4 is a closed-cell model (i.e., with no Ca^2+^ transport across the PM), which again is incompatible with KO experiments that change the total Ca^2+^ in the cell. Model 6 is unable to simulate MCU/NCLX-KO due to resultant instabilities in the model. The reason for this is the subject of current investigation; until we know why this happens any MCU/NCLX-KO simulations from Model 6 are unreliable.

**Figure 5.**
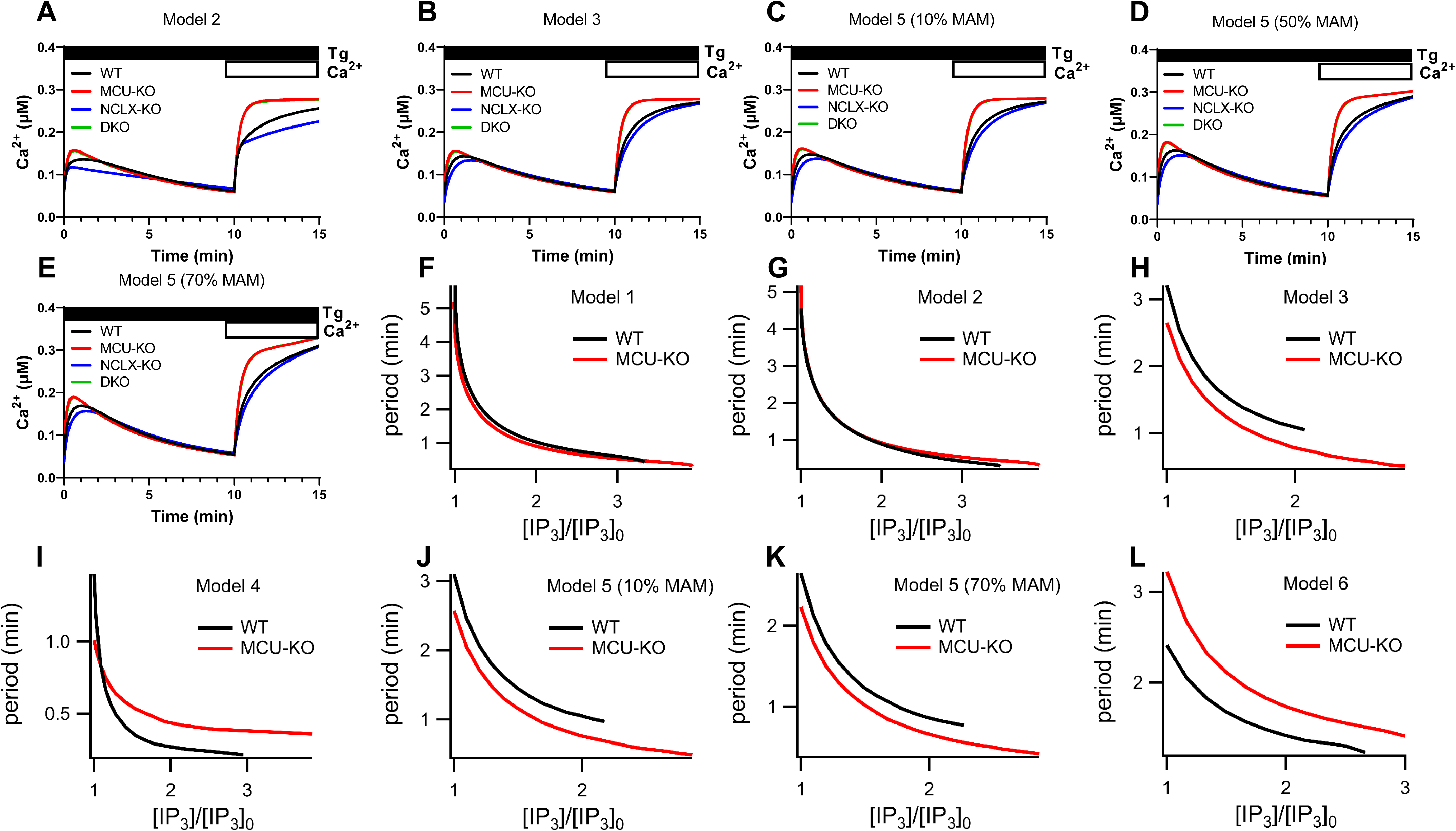
Mathematical modeling shows that MCU-KO increases cytosolic Ca^2+^ but has complex and non-linear effects on Ca^2+^ oscillations. (**A-E**), various models of the cytosolic Ca^2+^ signal in response to maximal store depletion with thapsigargin in the absence then presence of 2mM external Ca^2+^. In model 5, which considers MAMs regions of close contact between the ER and mitochondria, computations are shown when either 10%, 50% or 70% of total cellular IP_3_Rs are located within these MAMs. (**F-L**), Period of Ca^2+^ oscillations (1/Frequency) as a function of relative concentrations of IP_3_ ([IP_3_]_0_ is the lowest value of [IP_3_] for which oscillations exist in the model with MCU) for the six different models described in the Methods. For model 5, computations are shown when either 10% or 70% of total cellular IP_3_Rs are located within the MAMs.

When considering agonist-evoked Ca^2+^ oscillations, the qualitative results from each model differ significantly (**Fig. 5F-L**). Knockout of the MCU results in oscillations that, in some models, are slightly faster but in others are slightly slower. In some cases (Models 2 and 4), the effect on oscillations changes as agonist stimulation increases. Furthermore, in all cases, for a small range of even higher agonist concentrations, MCU-KO can turn a plateau response into an oscillatory response (this is the region where the “MCU-KO” curves overlap the wildtype “WT” curves). It follows that mitochondrial Ca^2+^ transport does not have a simple, monotonic, effect on the properties of Ca^2+^ oscillations. Instead, the effects of mitochondrial Ca^2+^ transport are critically dependent on the exact details of each mitochondrial Ca^2+^ flux, as well as other cellular parameters, such as the choice of IP_3_R model, the relative mitochondrial volume or the proportion of the ER membrane that is closely associated with the mitochondrial membrane.

Next, we constructed a model that considers MCU activity within a functional I_CRAC_ microdomain (IM model) whereby MCU activity prevents slow CDI of I_CRAC_. This model also considers that 10% of IP_3_Rs are within MAMs but does not include the equations for mitochondrial metabolism and membrane potential. The inclusion of the mitochondrial metabolism equations makes no qualitative difference to the results. Thus, the IM model is just Model 2 with the inclusion of I_CRAC_ CDI and MAMs with all other parameters being unchanged from Model 2. We modelled CDI of I_CRAC_ after(*32*) and initial tests of this IM model gave results that were qualitatively consistent with (*32*) (**Fig. S15**). We then simulated the effects of MCU-KO, NCLX-NO and MCU-KO + NCLX-KO on thapsigargin-activated cytosolic Ca^2+^ rise and agonist-evoked Ca^2+^ oscillations with three variations of this model that consider either 10%, 36% or 70% of the incoming Ca^2+^ through I_CRAC_ enters mitochondria (**Fig. S16**). The model showed that NCLX-KO leads to a slight decrease in SOCE activated by thapsigargin while MCU-KO and MCU-KO + NCLX-KO cause a similar increase in SOCE that recedes to a lower plateau equivalent to that of wildtype cells. Ca^2+^ oscillations in wildtype and MCU-KO cells in response to increasing concentrations of IP_3_ were not altered by either 10%, 36% or 70% variations in the IM model (**Fig. S16**).

In summary, our mathematical modeling predicts that all the experimental results presented here are consistent with the hypothesis that the MCU affects SOCE activated by thapsigargin and Ca^2+^ oscillations evoked by IP_3_-producing agonists only *via* its effect on mitochondrial Ca^2+^ transport, and that no microdomains connected to CRAC channels or IP_3_R (MAMs) are required to explain the experimental data.

## Discussion

Through Ca^2+^ uptake and extrusion, mitochondria can shape receptor-activated intracellular Ca^2+^ signaling. Mitochondria rapidly take up Ca^2+^ during receptor-evoked Ca^2+^ signaling and then extrude it much more slowly (*47*). Mitochondrial Ca^2+^ regulates metabolic activity, through modulation of the activity of three dehydrogenases of the TCA cycle(*48, 49*). Mitochondria can take-up cytosolic Ca^2+^ within the mitochondria-associated membrane (MAM) regions of close apposition between IP_3_R in the ER and mitochondria (*8*). In addition, respiring mitochondria can buffer Ca^2+^ from a distant site functionally connected to CRAC channels, thus reducing the extent of slow CDI of these channels to sustain SOCE and cytosolic Ca^2+^ signaling(*36, 50*).

In this study, we performed genetic knockout of the MCU in several non-excitable cell lines and in mice to determine the function of MCU in regulating receptor-evoked Ca^2+^ signaling. We reveal that MCU-KO enhances cytosolic Ca^2+^ signaling and downstream activation of NFAT transcription factors. Our data show that store depletion-mediated activation of SOCE and agonist-evoked cytosolic Ca^2+^ oscillations are enhanced when MCU-mediated mitochondrial Ca^2+^ uptake is abrogated. However, this enhancement of cytosolic Ca^2+^ signals is not due to increased activities of either IP_3_R or CRAC channels. We measured IP_3_R Ca^2+^ puffs in intact MCU-KO and wildtype control cells. The number of IP_3_R-mediated Ca^2+^ puffs and number of puff sites were similar between MCU-KO and wildtype cells. We recorded whole-cell CRAC currents from two cell lines under low physiological cytosolic Ca^2+^ buffering conditions with energized mitochondria and under strong cytosolic Ca^2+^ buffering and show that CRAC currents are of similar density in MCU-KO and wildtype parental controls. However, evaluation of CRAC channel slow CDI in the presence of thapsigargin showed that MCU activity prevents store-independent slow CDI of I_CRAC_, in agreement with previous studies(*34, 35*). Our data argue that MCU does not alter the activity of native IP_3_R. Although MCU function can affect CRAC channel activity by preventing its slow CDI and can inhibit cytosolic Ca^2+^ extrusion, the net result of MCU-KO is an enhancement of cytosolic Ca^2+^ after store depletion or agonist stimulation compared to control cells. Our findings have a simple and straightforward interpretation: abrogation of MCU-mediated Ca^2+^ uptake in response to receptor stimulation prevents the transfer of incoming Ca^2+^ through CRAC channels from the cytosol to mitochondria, causing cytosolic Ca^2+^ accumulation, increased ER Ca^2+^ refilling and enhanced NFAT nuclear translocation.

Using TIRF imaging, Korzeniowski et al. showed that on store depletion, mitochondria were remotely distant, or *ad minimum* excluded, from the sites of SOCE, which were visualized by STIM1 puncta and that mitochondrial Ca^2+^ uptake was independent of the distance between STIM1 and mitochondria(*51*). Naghdi et al. used as mAKAP-RFP-CAAX construct to immobilize mitochondria by linking them to the plasma membrane of endothelial cells and showed that this strategy has no effect on SOCE(*52*). Nevertheless, slow CDI of CRAC channels is mediated by a site that is hundreds of nm away, consistent with the reported localization of mitochondria within ∼200nm from the plasma membrane(*53*). Previous studies predating the discovery of MCU showed that abrogation of mitochondrial Ca^2+^ buffering using drugs such as the potent uncoupler of oxidative phosphorylation, FCCP or the inhibitor of Complex III of the ETC, Antimycin A1 inhibited SOCE(*35*). This decrease in SOCE resulted, at least partially, from evoking CRAC channel slow CDI(*34*). Our own findings revealed that mitochondrial depolarization with FCCP indeed inhibits SOCE, in agreement with these studies. However, FCCP equally inhibited SOCE in MCU-KO and wildtype cells, suggesting that although FCCP promotes I_CRAC_ slow CDI, the drug likely inhibits CRAC channels through additional MCU-independent mechanisms. Because these drugs are known to generate hydrogen peroxide(*54, 55*), their inhibitory effect on SOCE might be mediated by oxidation of Orai1(*46, 56*).

Extensive studies by Hoth and coworkers showed that mitochondria are critical for T-cell activation. Upon stimulation, mitochondria move closer, within 200 nm, of the immune synapse (IS) to sequester Ca^2+^. The presence of mitochondria near the IS coincides with enhanced Ca^2+^ concentration near the PM and inhibited cytosolic Ca^2+^ export through the PM Ca^2+^ ATPase (PMCA), thus enhancing the efficiency of NFAT activity and T-cell activation(*37, 53, 57, 58*). Therefore, both MCU function and its deletion can enhance NFAT activity. The former through inhibition of cytosolic Ca^2+^ export and CRAC channel slow CDI at the PM while the latter through enhanced global cytosolic Ca^2+^. Samanta et al. showed blunted SOCE and cytosolic Ca^2+^ oscillations in RBL-1 cells treated with siRNA against MCU and suggested that cytosolic Ca^2+^ uptake by MCU prevents CDI of both IP_3_R and CRAC channels(*36*). This discrepancy cannot be explained by differences between the MCU knockdown performed by Samanta et al. and MCU knockout in our study. We obtained similar results in primary splenic CD4^+^ T-cells and B-cells isolated from mice subjected to LoxP/Cre-mediated knockdown, which confers ∼60-80% MCU knockdown as documented by Westerns on isolated CD4^+^ T- and B-cells. Samanta et al. also reported that NCLX knockdown and Na^+^ depletion have no effect on CRAC channels(*50*), results that are at odds with previous findings (*35, 46*). Therefore, the reasons for the discrepancy between the study by Samanta et al. and ours are not clear. However, these investigators reported results from a single cell line, used one single siRNA, did not document protein or mRNA knockdown of MCU or NCLX and did not rule out off-target effects on STIM/Orai expression in their system(*36, 50*).

Our mathematical modeling support previous computational work showing that the effects of mitochondrial Ca^2+^ transport on cytosolic Ca^2+^ oscillations are complex and non-intuitive(*59*). Although mitochondria take-up and release cytosolic Ca^2+^ in a manner superficially similar to that of a traditional Ca^2+^ buffer, the complexities and nonlinearities inherent to these transport processes mean that the downstream effects on oscillation frequency cannot be easily predicted. In some cases oscillation frequency is increased by mitochondrial transport, in other cases oscillation frequency is decreased. Thus, the common understanding that inhibition of mitochondrial Ca^2+^ buffering decreases oscillation frequency is not supported by our modeling. Our models showed that MCU-KO enhances, while NCLX-KO decreases, cytosolic Ca^2+^ rise in response to store depletion by thapsigargin. Our modeling predicts that MCU-KO mediates its effects only through changes in mitochondrial Ca^2+^ transport, and that no Ca^2+^ microdomains that can alter CDI of Ca^2+^ channels (CRAC or IP_3_R) are required to explain our experimental data. Ishii et al showed that during Ca^2+^ oscillations in Hela cells Ca^2+^ shuttles between the ER and mitochondria and that Ca^2+^ uptake by mitochondria occurs at the expense of re-filling of the ER, in agreement with our observations(*60*). Our model of NCLX-KO is also consistent with previous experimental findings from our group and others. Hoth et al. showed that SOCE was reduced when NCLX was inhibited in cells subjected to Na^+^ depletion(*35*) and Naghdi et al. showed reduction in SOCE when cells were treated with CGP37157, a pharmacological inhibitor of the NCLX(*52*). We previously reported that molecular knockdown of NCLX or Na^+^ depletion inhibits SOCE and CRAC channel activity in HEK293 cells and primary vascular smooth muscle cells(*46*).

In summary, in addition to regulating CRAC channel inactivation, mitochondrial Ca^2+^ uptake by MCU is critical for interorganellar Ca^2+^ cycling and homeostasis, and its inhibition leads to enhanced cytosolic Ca^2+^ and NFAT induction.

## Materials and Methods

### Cell culture

Parental HEK293, Jurkat E6-1, RBL-1, DLD1, HCT116 and A20 cell lines were purchased directly from ATCC (Catalog # CRL-1573, TIB-152, CRL-1378, CCL-221,CCL-247, and TIB-208). HEK293, RBL-1, and HeLa cells were cultured in high glucose (4.5 g/L) Dulbecco’s modified Eagle’s medium (DMEM) supplemented with 10% heat-inactivated fetal bovine serum and 1X Antibiotic–Antimycotic solution. Jurkat, A20, and DLD1 were maintained in RPMI 1640 with L-glutamine supplemented with 10% heat-inactivated fetal bovine serum and 1X Antibiotic–Antimycotic solution. HCT116 were cultured in McCoys 5A medium supplemented with 10% heat-inactivated fetal bovine serum and 1X Antibiotic–Antimycotic solution. All cell lines were stored in a heated (37°C) humidified incubator under standard cell culture conditions (5% CO_2_; 95% air). The absence of Mycoplasma contamination was verified for all cell lines using a sensitive PCR based detection kit (ABM: G238).

### Fluorescence imaging

Adherent and semi-adherent cell lines were seeded onto round 25 mm glass coverslips twenty-four hours prior to imaging at a concentration of 1.5 x 10^6^ and 2.5 x 10^6^ cells respectively. Alternatively, A20, Jurkat E6-1, and primary CD4^+^ T cells were seeded onto round Poly-L-lysine (0.01%; mol wt 150,000-300,000) treated 25 mm glass coverslips 30 min prior to dye loading. Once attached, HEK293, HeLa, HCT116, and DLD1 cells were incubated in DMEM containing the ratiometric Ca^2+^ indicator Fura-2 AM (2 µM) for 30 min. RBL-1 cells were loaded with 4µM Fura-2AM following the same procedure. Jurkat and A20 cells were incubated with 2 µM Fura-2 AM in Hepes-buffered salt solution (HBSS; 120 mM NaCl, 5.4 mM KCl, 0.8 mM MgCl_2_, 20 mM Hepes, 10 mM Glucose adjusted to pH 7.4 with NaOH) supplemented with 2 mM Ca^2+^ for 30 min at room temperature. Before imaging, all coverslips were transferred from a standard 6-well plate into an Attofluor cell chamber and washed 3X with Ca^2+^ containing HBSS to remove excess Fura-2 AM. Using a fast shutter wheel (Sutter Instruments), cytosolic Ca^2+^ was measured by alternatively exciting Fura-2 with 340 nm and 380 nm wavelengths and recording the resulting 510 nm dye emission on a pixel by pixel basis through a 20X fluorescence objective paired with a Hamamatsu Flash 4 camera and processed using Leica Application Suite X software.

Measurement of NFAT4 nuclear translocation in HEK293 cells was achieved by transient overexpression of NFAT4-GFP (Addgene #21664) as previously described(*30, 40, 41*). Briefly, 1×10^6^ cells were incubated with a solution containing 1 µg of NFAT4-GFP and 3 µL Lipofectamine 2000 for 3 hrs. One day later, coverslips containing transfected cells were transferred to imaging chambers and washed with Ca^2+^ containing HBSS as described above. Using a Leica imaging system and a 40X oil immersion objective, NFAT4-GFP was excited at 488 nm and emission spectra selectively captured through a GFP filter cube. Real time analysis of NFAT4-GFP nuclear translocation in response to 10 µM Cch stimulation was calculated using the equation:

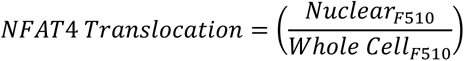

Quantification of ER Ca^2+^ store depletion and refilling was measured by transfecting parental and MCU-KO HEK293 cells with red R-CEPIA1er (Addgene: #58216) using Lipofectamine 24 hrs prior to imaging. Cells expressing R-CEPIA1er were then excited at 552nm and relative ER Ca^2+^ measurements recorded through a 40X oil immersion objective. Immediately after identifying R-CEPIA1er positive cells the bath solution was replaced with nominally Ca^2+^ free HBSS. One minute into the experiment cells were treated with CCh followed by washout and re-addition of 2 mM Ca^2+^ to the bath. The resulting traces represent ±SEM of all cells from at least three independent experiments.

### Ca^2+^ puff measurements using TIRF microscopy

Parental WT-HEK293 cells or MCU-KO cells were grown on 15-mm glass coverslips coated with poly-D-lysine (100 μg/ml) in a 35-mm dish for 2 days and imaged as previously described (*40*). Prior to imaging, the cells were washed three times with imaging buffer (137 mM NaCl, 5.5 mM glucose, 0.56 mM MgCl_2_, 4.7 mM KCl, 1.26 mM CaCl_2_, 10 mM HEPES, 1 mM Na_2_HPO_4_ at pH 7.4). Cells were subsequently incubated with Cal520-AM (5 µM; AAT Bioquest #21130) and 6-*O*-[(4,5-Dimethoxy-2-nitrophenyl)methyl]-2,3-*O*-(1-methylethylidene)-D-*myo*-Inositol 1,4,5-tris[bis[(1-oxopropoxy)methyl]phosphate] (ci-IP_3_/PM; 1 μM, Tocris #6210) in imaging buffer with 0.01 % BSA in dark at room temperature. 1 hour later, cells were washed three times with imaging buffer and incubated in imaging buffer containing EGTA-AM (5 μM, Invitrogen #E1219). After 45 minutes incubation, the media was replaced with fresh imaging buffer and incubated for additional 30 minutes at room temperature to allow for de-esterification of loaded reagents(*61*).

Following loading, the coverslip was mounted on chambers and imaged using an Olympus IX83 inverted total internal reflection fluorescence microscopy (TIRFM) equipped with oil-immersion Olympus UPLAAPO60XOHR (NA= 1.5) objective. The cells were illuminated using a 488 nm laser to excite Cal-520 and the emitted fluorescence was collected through a band-pass filter by a Hamamatsu ORCA-Fusion CMOS camera. The angle of the excitation beam was adjusted to achieve TIRF with a penetration depth of ∼140 nm. Images were captured from a field of view by directly streaming into RAM. To photorelease IP_3_, UV light from a 405 nm laser at 50 % power was introduced to uniformly illuminate the field of view. Both the intensity of the UV flash (∼2.4 mW) and the duration (1000 msec) for uncaging IP_3_ were optimized to prevent spontaneous puff activity in the absence of loaded ci-IP_3_. TIRF images were captured using 4 X 4 pixel binning (433.333 nm/pixel) from equal field of views for WT-HEK293 and MCU-KO cells at a rate of ∼166 frames per second. After visualizing images with the cellSens [Ver.2.3] life science imaging software (Olympus), images were exported as vsi files. Images, 10 seconds before and 60 seconds after flash photolysis of ci-IP_3_, were captured.

The vsi files were converted to tif files using Fiji and further processed using FLIKA, a Python programming based tool for image processing(*62*). From each recording, 500 frames (∼3 seconds) before photolysis of ci-IP_3_ were averaged to obtain a ratio image stack (F/Fo) and standard deviation for each pixel for recording up to 20 seconds following photolysis. The image stack was Gaussian-filtered, and pixel that exceeded a critical value (1.0 for our analysis) were located. The ‘Detect-puffs’ plug-in was utilized to detect the number of clusters (puff sites), number of events (number of puffs), amplitudes and durations of localized Ca^2+^ signals from individual cells. All the puffs identified automatically by the algorithm were manually confirmed before analysis(*63, 64*). The results from FLIKA were saved as excel and graphs were plotted using GraphPad Prism8.

### Patch clamp electrophysiology

RBL-1 cells were seeded onto 30 mm round glass coverslips 12 hours before recording; Jurkat cells were seeded onto Poly-L-lysine-coated coverslips 1 hour before recording. Native I_CRAC_ recordings were carried out using an Axopatch 200B and Digidata 1440A (Molecular Devices, LLC, Sunnyvale, CA) as previously described (Gonzalez-Cobos et al, 2013; Zhang et al, 2013, 2014). Pipettes were pulled from borosilicate glass capillaries (World Precision Instruments, Sarasota, FL) with a P-100 flaming/brown micropipette puller (Sutter Instrument Company, Novato, CA) and polished with DMF1000 (World Precision Instruments, Sarasota, FL). Resistances of filled glass pipettes were 2–4 MΩ. Only cells with tight seals (> 16 GΩ) were selected for break-in. Cells were maintained at a +30 mV holding potential during experiments and subjected to a 250 ms voltage ramps from 100 mV to -140 mV every 3 seconds. Reverse ramps were designed to inhibit Na^+^ channels potentially expressed in these cells. All experiments were performed at room temperature. 8 mM MgCl_2_ was included in the pipette solution to inhibit TRPM7 currents. CRAC currents were induced by either 30 µM inositol-1,4,5-trisphosphate (1,4,5-IP_3_) or 50 mM BAPTA. Clampfit 10.3 software was used for data analysis. The solutions employed for patch-clamp recordings are as follows.

Bath solution:

115 mM Na-methanesulfonate, 10 mM CsCl, 1.2 mM MgSO_4_, 10 mM Hepes, 20 mM CaCl_2_, and 10 mM glucose (pH 7.4, adjusted with NaOH).

Divalent-Free (DVF) Bath Solution:

155 mM Na-methanesulfonate, 10 mM HEDTA, 1 mM EDTA, and 10 mM HEPES (pH 7.4, adjusted with NaOH).

Pipette solution 1 (Low Buffer):

135 mM Cs-methanesulfonate, 1 mM EGTA, 8 mM MgCl_2_, and 10 mM HEPES, 0.03 mM IP_3_, 2 mM pyruvic acid, 2 mM malic acid, 1 mM NaH_2_PO_4_, 2 mM Mg-ATP (pH 7.2 adjusted with CsOH)

Pipette solution 2 (High Buffer):

85 mM Cs-methanesulfonate, 50 mM Cs-1,2-bis-(2-aminophenoxy)ethane-N,N,N′,N′-tetraacetic acid (Cs-BAPTA), 8 mM MgCl_2_, and 10 mM HEPES (pH adjusted to 7.2 with CsOH).

Recordings of CRAC channel store-independent slow CDI were performed according to(*34*). Cells were incubated in the bath solution without Ca^2+^ and in the presence of 2 µM thapsigargin for 3-5 min before beginning of recordings. The bath and pipette solutions were as follow:

Bath solutions:

0 mM Ca^2+^ bath solution (in mM): 155 NaCl, 4.5 KCl, 3 MgCl_2_, 10 D-glucose, and 5 Hepes (pH=7.4 with NaOH)

20 mM Ca^2+^ bath solution (in mM): 125 NaCl, 4.5 KCl, 20 CaCl_2_, 3 MgCl_2_, 10 D-glucose, and 5 Hepes (pH=7.4 with NaOH)

Pipette solutions:

Low buffer pipette solution (in mM): 140 Cs-methanesulfonate, 2 MgCl_2_, 0.66 CaCl_2_, 1.2 EGTA, 10 Hepes (pH=7.2 with CsOH)

Low buffer + mitochondrial cocktail pipette solution (in mM): 140 Cs-methanesulfonate, 2 MgCl_2_, 0.66 CaCl_2_, 1.2 EGTA, 10 Hepes, 2.5 mM malic acid, 2.5 mM pyruvate acid, 1 mM NaH_2_PO_4_, 5 mM Mg-ATP, 0.5 mM Na-GTP (pH=7.2 with CsOH)

### Cytosolic and mitochondrial volume determination

To determine the cytosolic volume of all cell lines pEGFP-N1 (Addgene# 6085-1) was transiently overexpressed. One day before cytosolic volume measurement all cell lines were electroporated using the Amaxa Nucleofector II. Cells were first resuspended in the transfection reagent supplemented with pEGFP-N1 (1 µg). Cell line specific protocols Q-001, X-001, T-030, and L-013 were used to electroporate HEK293, Jurkat, RBL-1, and A20 cells respectively. One day later, live cells were stained with MitoTracker Deep Red (200 nM) for 15 min at room temperature in the dark before imaging on a Leica DMI8 confocal microscope. Stained and washed cells were observed through a 40X oil immersion objective. Both GFP and MitoTracker signals were then collected from randomly selected fields of view by compiling Z-Stacks of the lower and upper bounds of all cells. MitoTracker and GFP channels were then used to reconstruct 3D surfaces using IMARIS Cell Biology software (Oxford Instruments). Once both cytosolic (GFP) and mitochondrial (MitoTracker) surfaces were reconstructed, volume measurements were recorded individually.

### ImageStream native NFAT1 nuclear translocation assay

Jurkat Cells were first counted and transferred to 1.5mL Eppendorf tubes at a concentration of 1.2×10^6^ cells per replicate. Cells were then centrifuged, and media replaced with HBSS supplemented with 2mM Ca^2+^ to eliminate the possibility of serum mediated activation. Experimental cells were then stimulated with α-CD3 (0.125µg/mL) or treated with Tg (2µM) to induce cytosolic Ca^2+^ oscillations or evoke SOCE. Following stimulation, cells were immediately fixed and permeabilized using a transcription factor staining kit according to the manufacturer protocol (Tonbo Biosciences Cat: TNB-0607-KIT). After fixation and permeabilization cells were spun down, the supernatant removed, and stained with a primary conjugated NFAT1-FITC antibody (Cell Signaling #14324S @ 1:100) for 1hr at room temperature protected from light. Unbound NFAT1 antibody was then washed away and cells resuspended in 40uL of PBS. Nuclear staining was performed immediately before image acquisition by adding 10uL of the nuclear stain Dapi (5X) to give a final concentration of (0.25µg/mL).

Images were acquired using a 10-color Amnis ImageStream X Mk II imaging cytometer. Channels 1 and 9 (430-480nm) were used to obtain bright field images, channel 2 for FITC (Ex:488 Em:480-560nm), and Dapi through channel 7 (Ex:405 Em:430-505nm). Prior to final image acquisition cells were gated to exclude debris and doublets and Ex laser power adjusted to prevent pixel saturation. From each stained and unstained sample 3000-5000 in focus single-cell events were collected. After running all samples single color compensation samples were run for FITC then Dapi. All data was analyzed using IDEAS software version 6.2 (Amnis Corp, Seattle, WA). During the analysis in focus cells were first identified, then doublets excluded, and lastly cells positive for both DAPI and NFAT1 were selected for further analysis. The extent of nuclear NFAT1 was then determined using the nuclear translocation module. Raw nuclear similarity scores were then exported to Prism GraphPad resulting in the final figures.

### Generation of transgenic MCU^fl/fl^ CD4^Cre(Ert2)^ and MCU^fl/fl^ MB1^Cre^ mice

We generated inducible MCU-KO CD4^+^ T cells by breeding MCU^flx/flx^ mice (generous gift from Dr. John W. Elrod, temple University)(*65*), with CD4CreER^T2^ (B6(129X1)-Tg(Cd4-cre/ERT2)11Gnri/J) purchased from Jackson labs(*66*). Resulting CD4CreER^T2^ positive MCU^fl/wt^ mice from this cross were then bred with MCU^fl/fl^ mice to generate all control and experimental cohorts. B-cell specific knockout of MCU was achieved by breeding MB1-Cre (B6.C(Cg)-Cd79a^tm1(cre)Reth^/EhobJ) mice from Jackson laboratory with MCU^fl/fl^ mice. From that cross MCU^fl/wt^MB1Cre^+/-^ were then bred with MCU^fl/fl^ mice to generate the experimental and control groups (*67*). PCR based genotyping was performed on all mice to identify experimental and control mice. Western blots were used to validate MCU-KO at the protein level. Both male and female mice were utilized in equal numbers to reduce the possibility of sex dependent variation biasing the results. All mice were maintained in the specific pathogen free barrier animal facility at the Pennsylvania State University College of Medicine animal facility. Prior to any breeding or experimentation all protocols used were approved by the Pennsylvania State Institutional Animal Care and Research Advisory Committee (IACUC) to ensure all animals were handled and cared for according to ethical guidelines.

### Primary T- and B-cell isolation and culture

Six to eight-week-old MCU^fl/fl^ CD4^Cre+^ or MCU^fl/fl^ MB1^Cre+^ mice and control counterparts were humanly sacrificed immediately prior to immune cell isolation. Superficial cervical and Inguinal lymph nodes were first collected followed by the spleen. All lymphoid tissues were then disrupted by passing the tissue through a 70 µm mesh screen. The resulting single cell suspension was than washed with 10 mL of cold RPMI and resuspended in 1 mL of Robosep buffer (STEMCELL technologies) prior to the negative selection of B- or T-cells. Using the corresponding negative selection kit primary mouse CD4^+^ T-cells and B-cells were isolated using the magnetic beads and antibody cocktail as recommended by the manufacturer.

Isolated CD4^+^ T-cells were then cultured *ex vivo* to increase the efficiency of CRE^ERT2^ activity. Naïve Primary CD4^+^ T were maintained in RPMI 1640 supplemented with 10% FBS, 1X Antibiotic-Antimycotic, 1X Glutamax, mIL2 (30 U/mL), and Mouse T-Activator anti-CD3/CD28 Dynabeads (Thermo Fisher) at a concentration of one bead per cell. Experimental mice (MCU^fl/fl^ CD4^Cre+^) and control (MCU^fl/fl^ CD4^Cre-^) mice were both treated with 2 µM 4-OHT for four days. As a control, the second group of (MCU^fl/fl^ CD4^Cre-^) mice was only administered vehicle (DMSO). Cells were subsequently washed and beads removed.

### Genetic knockout of MCU in multiple cell lines using CRISPR/Cas9

To generate MCU-KO HEK293 and HCT116 cells, human MCU gRNA3 (hMCU gRNA1) was subcloned into the lentiCRISPR v2 backbone (Addgene, Plasmid #52961) and transfected into wild-type HEK293 cells *via* nucleofection as previously described. Forty-eight hours after transfection, CRISPR/Cas9 positive cells were selected by treating the culture medium with puromycin (2 μg/ml) (Gemini Bio Products, West Sacramento, CA). After six-days of selection, cells were trypsinized and seeded at a concentration of 0.5/cells per well into a 96 well plate. Resulting colonies were then screened using the Guide-it Mutation Detection Kit (Clontech Laboratories, 631443). Knockout was further confirmed via western blot, functional analysis, and genomic sequencing (Fig. 1).

An alternative method was used for all other cell lines to avoid any potential effects of constitutive Cas9 expression. The same two human gRNAs (hMCU gRNA1and 2) were used to generate all Jurkat and DLD1 MCU-KO clones. Two independent mouse specific gRNAs (mMCU gRNA 1 and 2) were used to generate MCU-KO A20 cells. While a single gRNA (rMCU gRNA1) was used to generate MCU-KO RBL-1 cells. All gRNA sequences can be found in **supplementary table 1**. gRNAs for all cell lines were cloned into one of two fluorescent vectors to allow for the identification of transiently transfected cells using fluorescence-activated cell sorting (FACS) (pSpCas9(BB)-2A-GFP and pU6-(BbsI)_CBh-Cas9-T2A-mCherry: Addgene). One day after transfection with the corresponding gRNA single cells with high gRNA expression were sorted into the wells of a 96 well plate using a FACS Aria SORP high-performance cell sorter. After sorting cells were maintained in complete medium until colonies began to form ∼2-3 weeks depending on the cell line. Visible colonies were collected and analyzed by Western blot to identify knockout clones. Positive clones were functionally analyzed and those clones that showed no mitochondrial Ca^2+^ uptake were used for all experiments.

Furthermore, all clones were sequenced to validate gRNA efficiency. In HEK293 MCU-KO cells we observed two different genomic deletions (11nt and 1nt; exon 2) that resulted in a genomic frameshift and the formation or early stop codons. In Jurkat MCU-KO cells we also observed two different mutations. The first was a 7nt insertion into exon 3 and the second caused multiple indels that resulted in a genetic frameshift and formation of an early stop codon in exon 3. In RBL-1 we observed two different deletions from exon 3 (16 and 400nt) that both resulted in genomic frameshifts. Two different deletion events were also observed in A20 cells, the first was a single nucleotide deletion and the second a 7nt deletion from exon 1. We also observed two different deletions in HCT116. The first being a 16nt deletion and the second a 19nt deletion; both occurred in exon2 and resulted in frameshifts. In DLD1 cells, two unique 2nt deletions were observed within exon 4 that both resulted in genomic frameshifts and the introduction of an early stop codon.

### Western blot analysis

Cells were collected from culture flasks or isolated from mice, washed with ice cold phosphate-buffered saline, and pelleted via centrifugation prior to lysis in ice-cold RIPA buffer (150 mM NaCl, 1.0% IGEPAL CA-630, 0.5% sodium deoxycholate, 0.1% SDS, 50 mM Tris pH 8.0; Sigma) supplemented with protease and phosphatase inhibitor (Halt; Thermo Scientific) for 15 min on ice. Crude protein lysates were clarified by centrifugation (15,000 x g/12 min @ 4°C) and the supernatant collected. Total protein concentration was determined using the Pierce BCA assay as previously described prior to loading a NuPAGE precast 4-12% Bis-Tris gel. After subjecting the samples to 120V for 1.5 hrs proteins were transferred to a polyvinylidene difluoride membrane using a BioRad Criterion blotter and 1X NuPAGE Transfer Buffer (Invitrogen). After 1 hr of transferring at 95 V the membrane was removed and blocked with LI-COR Tris-buffered saline (TBS) buffer for 1 hr at room temperature. Primary antibodies were added to blocked membranes overnight at 4°C in a shaker. Nest day, PVDF membranes were then washed 3X with TBST and probed with the corresponding species-specific LI-COR secondary antibody for 1 hr at room temperature. Then, the membrane was washed 3X with TBST and immediately imaged using a LI-COR odyssey imaging system and fluorescence quantified using Image Studio lite software (LI-COR).

### Measurement of mtΨ and mtCa^2+^ uptake in permeabilized cells

HEK293 (6×10^6^), Jurkat (10×10^6^), A20 (10×10^6^), and RBL-1 (7×10^6^) cells were washed with PBS immediately before resuspension in an intracellular medium (ICM) consisting of 10mM NaCl, 120 mM KCl, 1 mM KH_2_PO_4_, 20 mM Hepes-Tris, pH 7.2 and 2 μM of the SERCA pump inhibitor thapsigargin. Just before beginning the measurement 2 mM succinate was added to the ICM. Cells were permeabilized with 40 µg/mL digitonin and simultaneous measurements of mitochondrial membrane potential (mtΨ) and extra-mitochondrial Ca^2+^ was achieved by loading permeabilized cells with the ratiometric mtΨ dye JC-1 (800 nM) and the fluorescent Ca^2+^ indicator Fura2-FF (0.5 µM). Both dyes were excited, and emissions recorded using a dual wavelength excitation and emission fluorimeter (Delta Ram, PTI).

### qRT-PCR analysis

Total mRNA was isolated from HEK293 (2×10^6^) and Jurkat cells (4×10^6^) parental and MCU-KO cell lines using the RNeasy Mini Kit (Qiagen). Isolated RNA was then analyzed using a NanoDrop spectrophotometer (Thermo Scientific) and 1 µg of DNAse I treated (Thermo Fisher) RNA was reverse transcribed using the High-Capacity cDNA Reverse Transcription Kit (Applied Biosystems). Total cDNA (1 µL) was diluted 1:5 then combined with target-specific primers, SYBR Green qPCR Master Mix (Applied Biosystems), and mqH_2_O resulting in a 10 µL total reaction. All targets were subjected to the same PCR protocol that began with an initial activation step for 2 min at 50 °C followed by a 95 °C for 2 min melt step. The initial melt steps were then followed by 40 cycles of 95 °C for 15 s, a 15 s annealing step at 54.3 °C, and target amplification at 72 °C for 30 s. Once complete, a standard melt curve was generated to ensure primer specificity. Analysis of target and control samples was carried out using the comparative Ct method. All samples were normalized to the average of the two reference genes GAPDH (glyceraldehyde-3-phosphate dehydrogenase) and NONO (Non-POU Domain Containing Octamer Binding). All samples were run in triplicate to ensure reproducibility.

### Mathematical Modeling

Models of the IP_3_R, SERCA and the PM ATPase were taken directly from(*40, 41*), with unchanged parameters. The model of Ca^2+^ influx, however, was simplified from that of (*40, 41*) by omitting the majority of the complexity generated by two different forms of STIM and three different forms of ORAI. Instead we modelled Ca^2+^ influx by

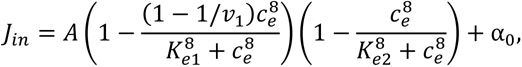

where A=0.2 µM/s, v_1_=3, K_e1_=50 µM, K_e2_=200 µM and α_0_=0.005 µM/s. The variable c_e_ denotes ER [Ca^2+^].

To incorporate mitochondrial transport, the base model was supplemented by the addition of three terms each representing a mitochondrial Ca^2+^ flux: a flux through the MCU (J_MCU_), a flux through the NCLX (J_NCLX_), and (following (*42*)) a background leak term (J_x_). The four different models of mitochondrial Ca^2+^ transport are as follows.

Model 1. The mitochondria are modelled as simple buffers. Thus

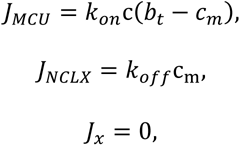

where k_on_=3 µM^-1^s^-1^, k_off_=5/s, b_t_=5 µM were chosen so as to give a physiological resting value of c_m_, and to ensure that the buffering was close to linear. The variable c_m_ denotes the mitochondrial [Ca^2+^].

Model 2. The mitochondrial fluxes are modelled following (*42*) but with the omission of the mitochondrial membrane potential and the mitochondrial metabolism. Thus

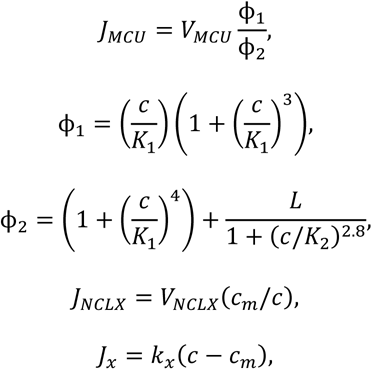

where V_MCU_ = 15 µM/s, K_1_=6 µM, K_2_=0.38 µM, V_NCLX_=0.005 µM/s, L=50 and k_x_ =0.01/s. The variable c denotes the cytosolic [Ca^2+^].

Model 3. The mitochondrial fluxes are modelled following (*42*) (and use the same parameter values), including the mitochondrial membrane potential and the equations describing mitochondrial metabolism. As with the previous models, the model of the IP_3_R is taken from(*23, 40, 41*), and thus the results from Model 3 are not identical to those of (*42*). No mitochondria-associated membrane (MAM) microdomains are included.

Model 4 is identical to the model of (*42*).

Model 5. The mitochondrial fluxes are identical to those of Model 3, but now MAMs are included, in the manner of (*45*) (i.e., using a compartmental model approach, and not including any spatially distributed compartments). However, the IP_3_R model is the same as in Model 3, and thus Models 5 and 6 give significantly different outcomes. To maintain a physiological resting c_m_, V_MCU_ was increased to 150 µM/s.

Model 6 is identical to the model of (*45*).

#### Model for calcium-dependent inactivation of I_CRAC_

A specified fraction (either 10%, 36% or 90%) of J_in_ was assumed to enter the cell directly into an I_CRAC_ microdomain (IM) formed between the PM and the mitochondrial membrane, containing only I_CRAC_ and MCU. Inclusion of NCLX and/or PM Ca^2+^ pumps into the IM makes no qualitative difference to the results. To describe calcium-dependent and time-dependent inactivation of I_CRAC_ we used one of the simplest possible models. A new time-dependent variable, h_icrac_ was introduced, obeying the differential equation

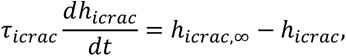

Where

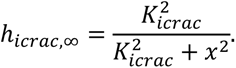

Here, x is either the cytosolic or IM [Ca^2+^] depending on whether we are modeling the influx into the IM or into the cytosol. Finally, J_in_ was replaced by h_icrac_J_in_, so that an increase in [Ca^2+^] at the IM causes inhibition of the CRAC channel, with time constant t_icrac_. Calcium moves from the IM to the cytosol at a rate proportional to the concentration difference of the two compartments, with rate constant 0.001/min. Other parameter values (K_icrac_=0.05 mM, t_icrac_=5 min) were chosen to give qualitative agreement with Fig. 2A of (*32*) (as shown in **Fig. S15**).

### Statistical analysis

All statistical analysis was performed using GraphPad Prism version 9. Statistically significant differences between groups were identified within the figures where *, **, ***, and **** indicate p-values of < 0.05, < 0.01, < 0.001, and <0.0001 respectively. All statistical tests and resulting p-values from that test are recorded within the figure legends.

## Acknowledgments

We are grateful to Dr. Priya Santhanam (Penn State University) for help and advice on Ca^2+^ measurements in permeabilized cells and for giving us unlimited access to her PTI imaging system, Dr. Suresh Joseph (Thomas Jefferson University) for the kind gift of HeLa cells and their MCU-KO clone, Dr. John W. Elrod (Temple University) for kindly providing us with the MCU^flx/flx^ mice, and Drs. Jeff Lock and Ian Parker (University of California, Irvine) for their advice and help with the Ca^2+^ puff measurements using TIRF microscopy and with data analysis. This work was supported by NIH/NHLBI (R35-HL150778 to MT), NIH/NHLBI (R01 HL137852 to Scott Earley and MT) NIH/NIDCR (R01 DE019245 to DIY and JS), NIH/NCI (R01 CA242021 to NH), and by the NIH/NIDDK Intramural Research Program (to JMH).

## Author contributions

REY, SME, DIY, JS and MT designed research; REY, SME, XZ, XP, VA, NL, TP, JCB and MTJ performed research; JMH, GD and JS conceived and performed mathematical simulations; DIY, and NH contributed new reagents/analytic tools; REY, SME, XZ, VA, DIY and MT analyzed data; and REY and MT wrote the paper with input from all authors.

## Data and materials availability

Statistical analyses and raw data of all experiments herein including unprocessed Westerns are included in a separate file. All original cell lines reported herein are available from the lead investigator upon request and completion of an MTA.

---

**Supplementary Figure 1.**
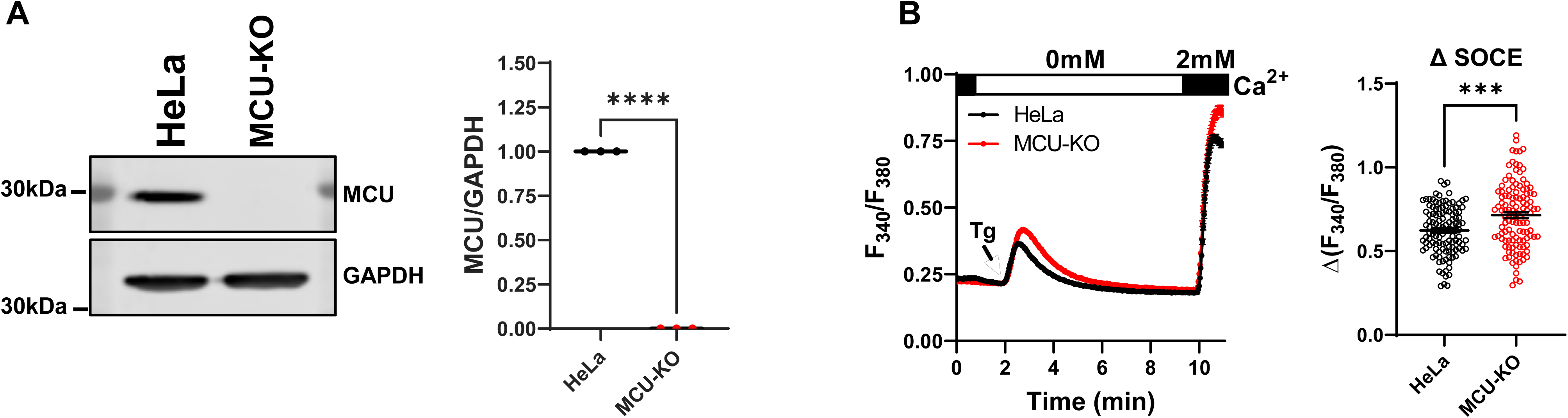
MCU-KO in HeLa cells enhances cytosolic Ca^2+^ upon thapsigargin stimulation. Western blot documenting MCU protein knockout in MCU-KO of HeLa cells compared to parental cells (**A**). Quantification of MCU protein band densitometry relative to GAPDH from three independent experiments are graphed (**A**). Ca^2+^ measurements in HeLa parental cells and MCU-KO cells in response to stimulation with 2 µM thapsigargin (Tg) in Ca^2+^-free buffer followed by restoration of 2 mM extracellular Ca^2+^ to determine the magnitude of SOCE. Quantification of SOCE (Δ SOCE) from at least three independent experiments are also shown (**B**).

**Supplementary Figure 2.**
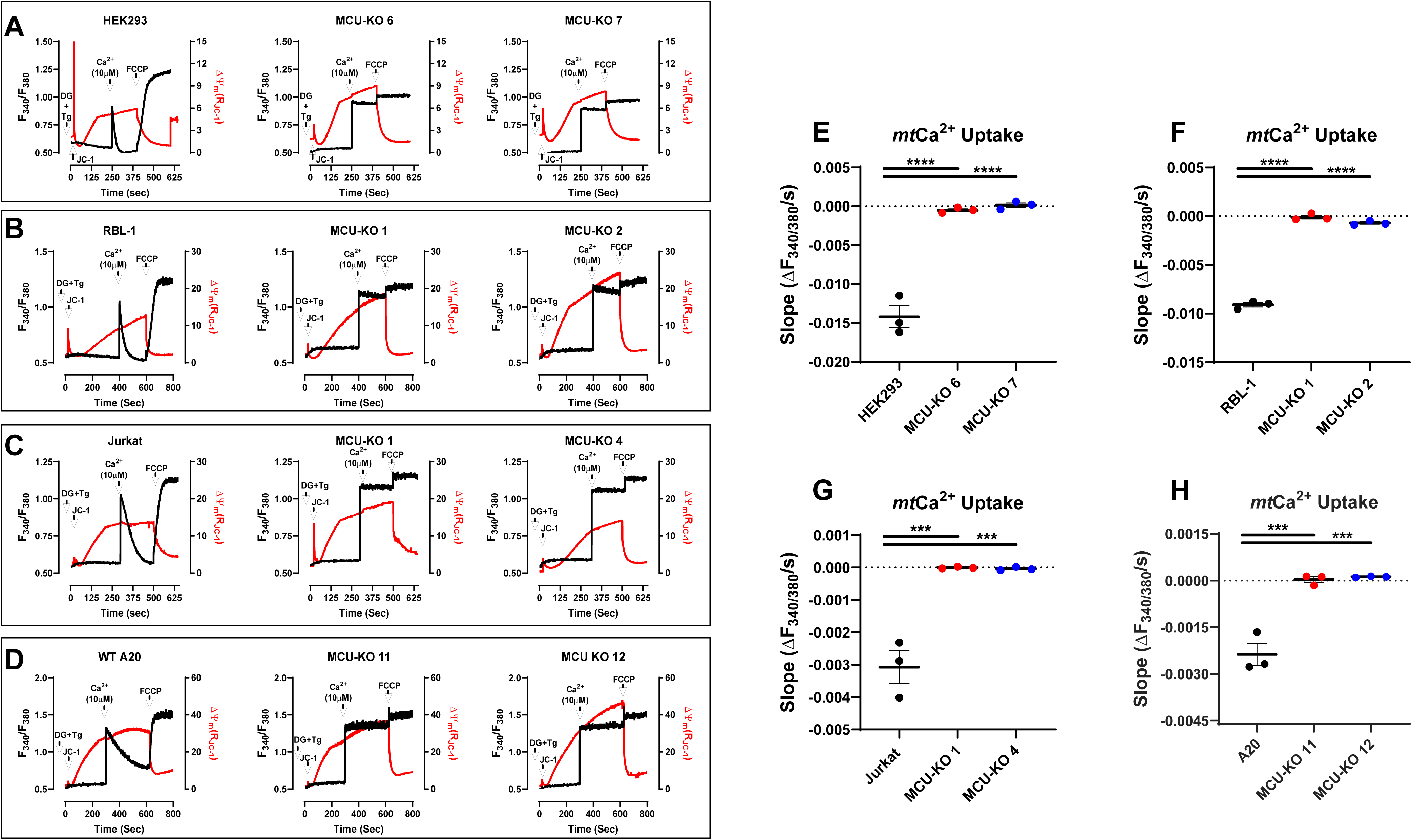
MCU-KO cells have abrogated mitochondrial Ca^2+^ uptake. **(A)**, Simultaneous measurements of mitochondrial Ca^2+^ uptake and membrane potential in permeabilized cell populations of wildtype and MCU-KO HEK293 cells. (**B-D**), Similar measurements to (**A**) in RBL-1, Jurkat and A20 cells, respectively. (**E-H**), Quantification of mitochondrial Ca^2+^ uptake in wildtype and MCU-KO of HEK293, RBL-1, Jurkat and A20 cells, respectively.

**Supplementary Figure 3.**
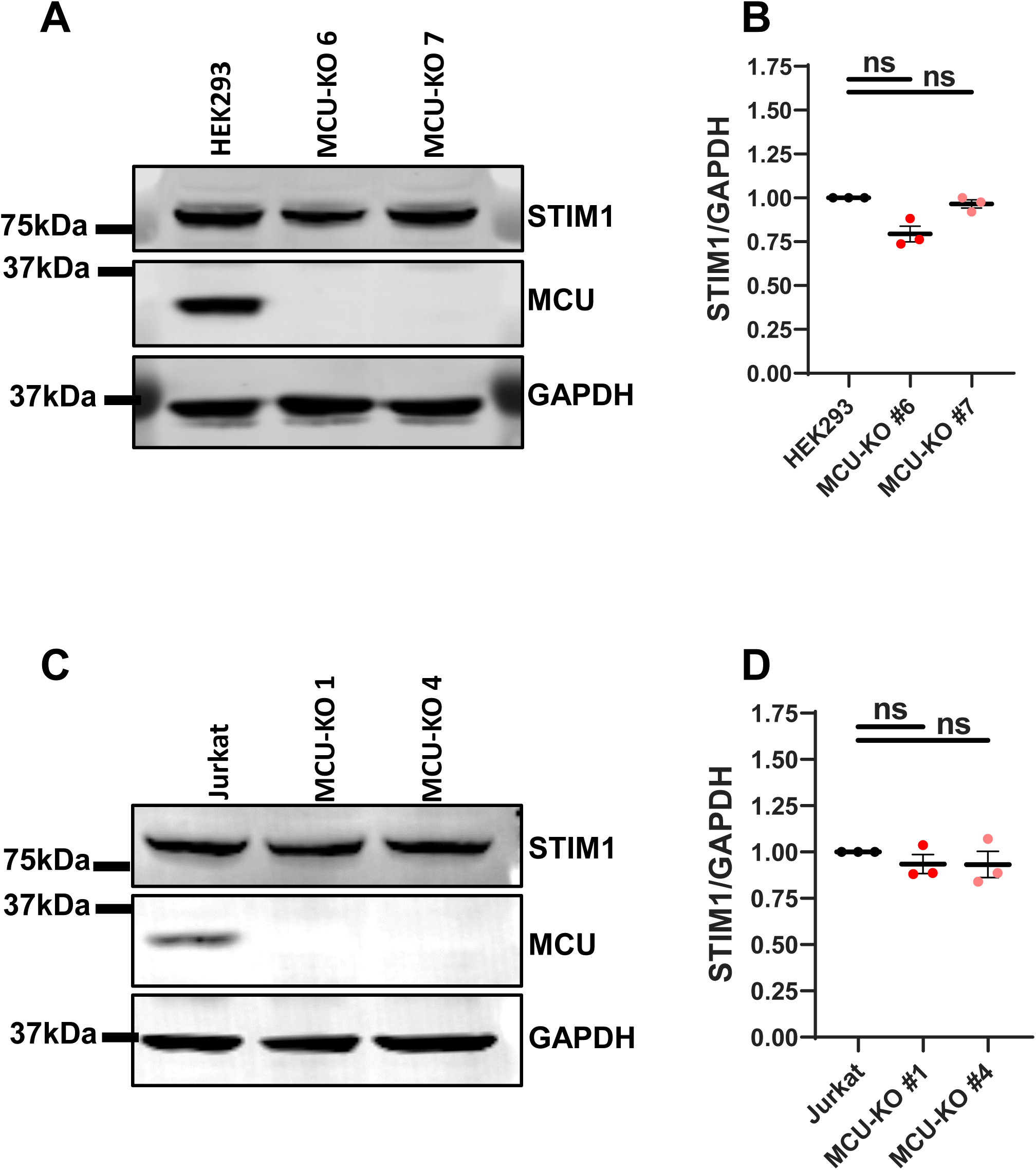
MCU-KO does not alter STIM1 protein expression. **(A)**, Western blot for STIM1 and MCU proteins in wildtype and MCU-KO HEK293 cells. (**B**) Quantification of STIM1 band density relative to GAPDH from three independent experiments similar to (**A**). (**C, D**), Similar experiments to (**A, B**) in wildtype and MCU-KO Jurkat cells.

**Supplementary Figure 4.**
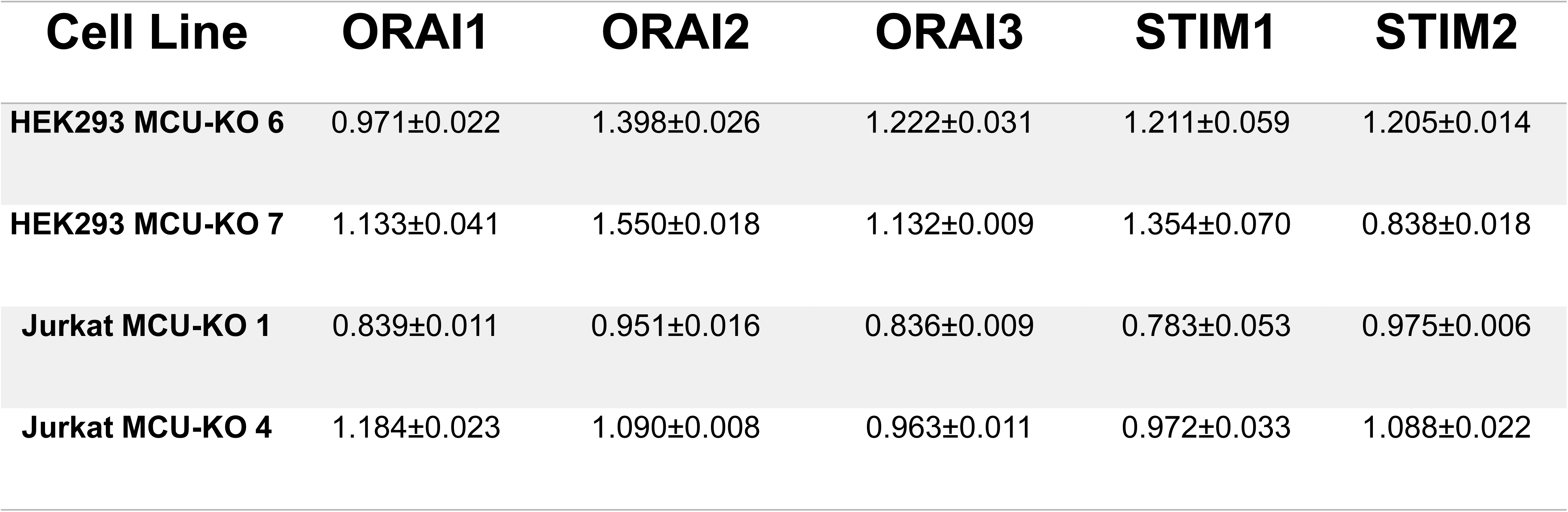
MCU-KO does not significantly affect mRNA expression of STIM/Orai isoforms. QPCR on STIM and Orai isoforms from wildtype and MCU-KO cells of HEK293 and Jurkat.

**Supplementary Figure 5.**
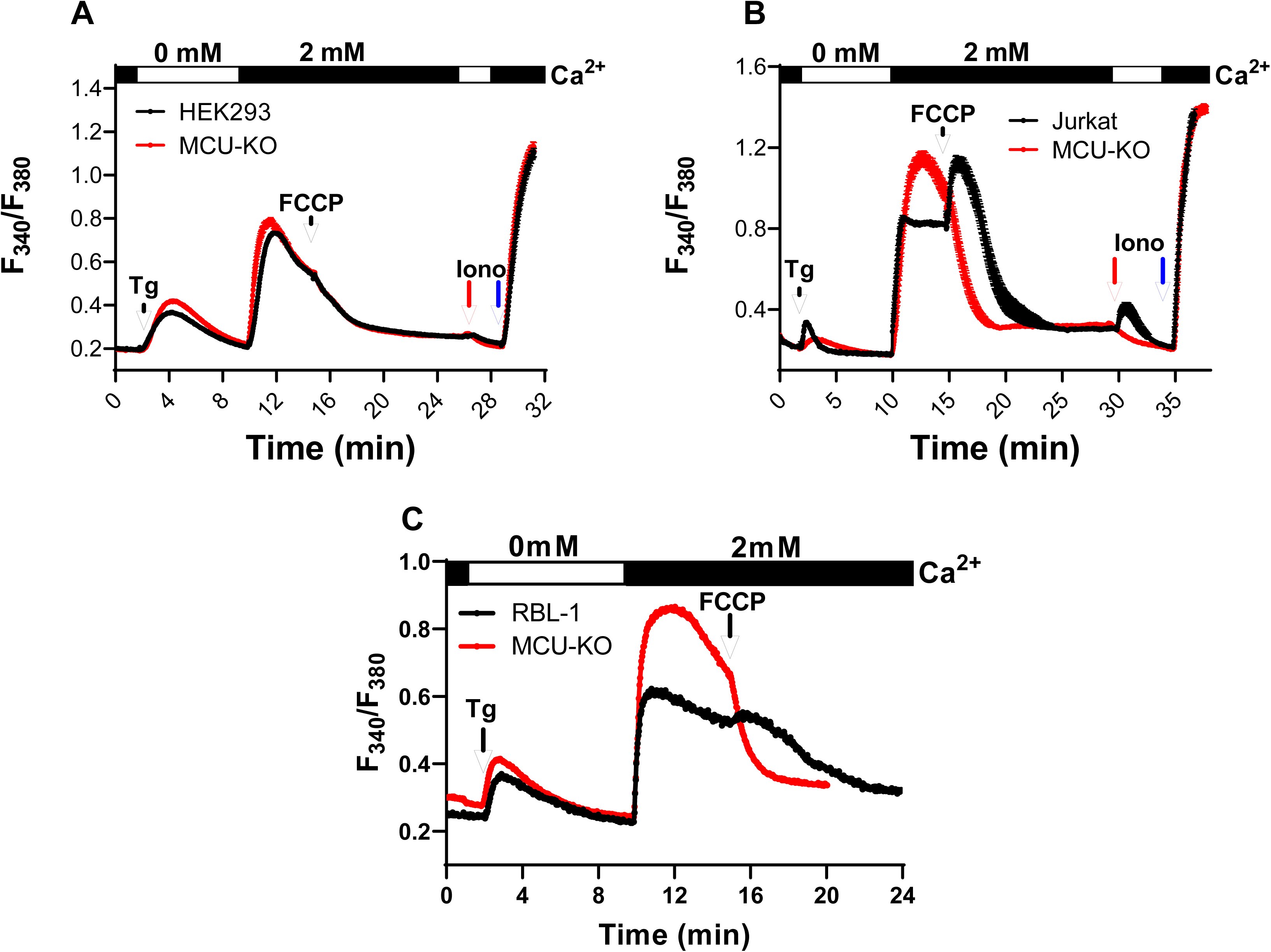
Dissipation mitochondrial membrane potential inhibits SOCE in wildtype and MCU-KO cells. **(A-C)**, Fura2 Ca^2+^ measurements using 2 µM thapsigargin first in the absence then presence of 2 mM extracellular Ca^2+^ in wildtype and MCU-KO HEK293 (**A**), Jurkat (**B**) and RBL-1 (**C**) cells. Addition of 5 µM FCCP to cells after Ca^2+^ entry is initiated, followed in (**A, B**) by 1µM ionomycin in 0 mM external Ca^2+^ (red arrow), then 10µM ionomycin in 2 mM external Ca^2+^ solution (blue arrow).

**Supplementary Figure 6.**
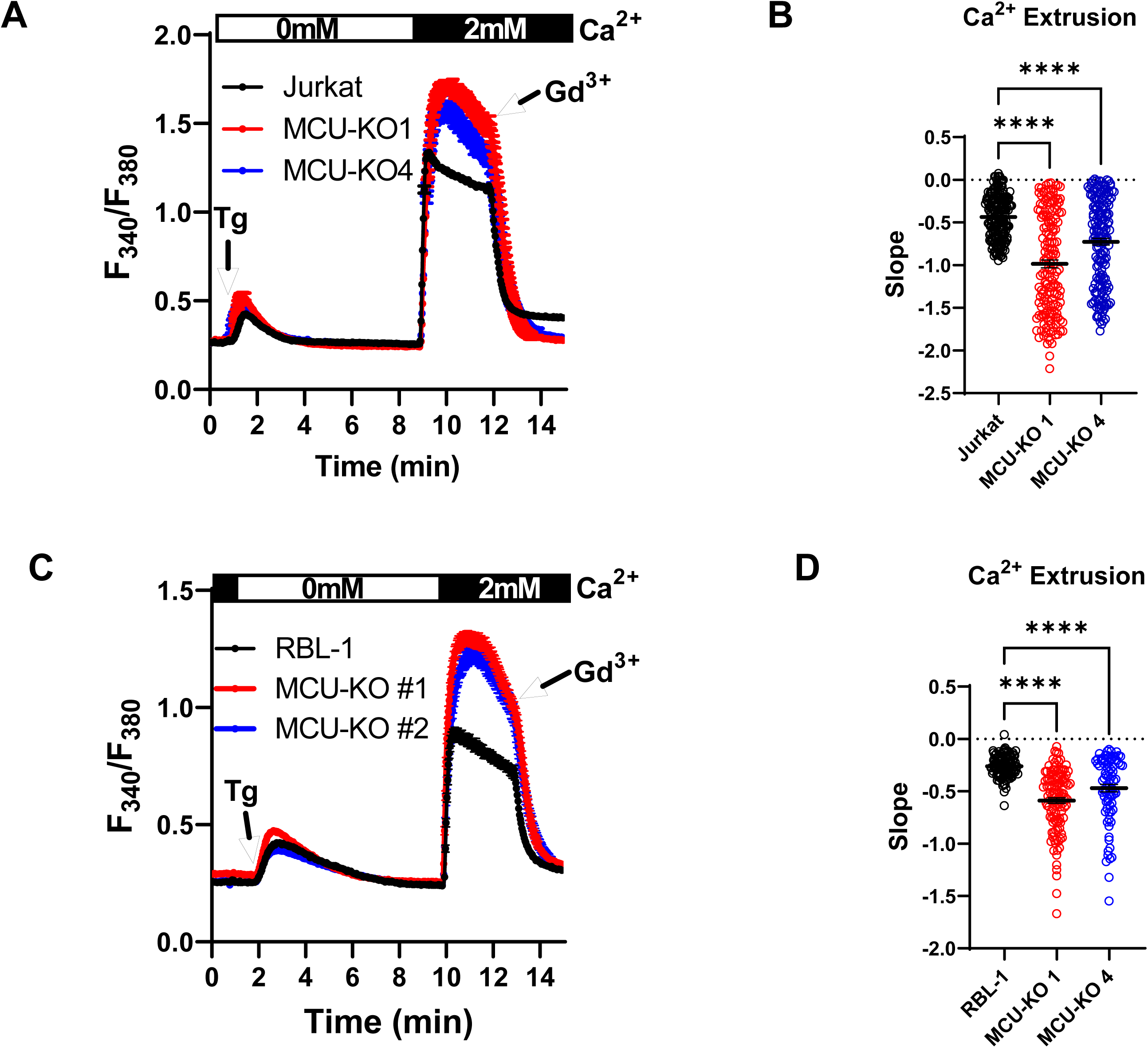
MCU-KO enhances Ca^2+^ extrusion. MCU-KO and parental Jurkat (A, B) and RBL-1 (C, D) cells were stimulated with 2 µM thapsigargin (Tg) in Ca^2+^-free buffer followed by restoration of 2 mM extracellular Ca^2+^ to determine the magnitude of SOCE followed by addition of 5 µM Gd^3+^ to block SOCE and assess Ca^2+^ extrusion. The slope of Ca^2+^ extrusion in these cells was calculated and reported statistically in (**B, D**).

**Supplementary Figure 7.**
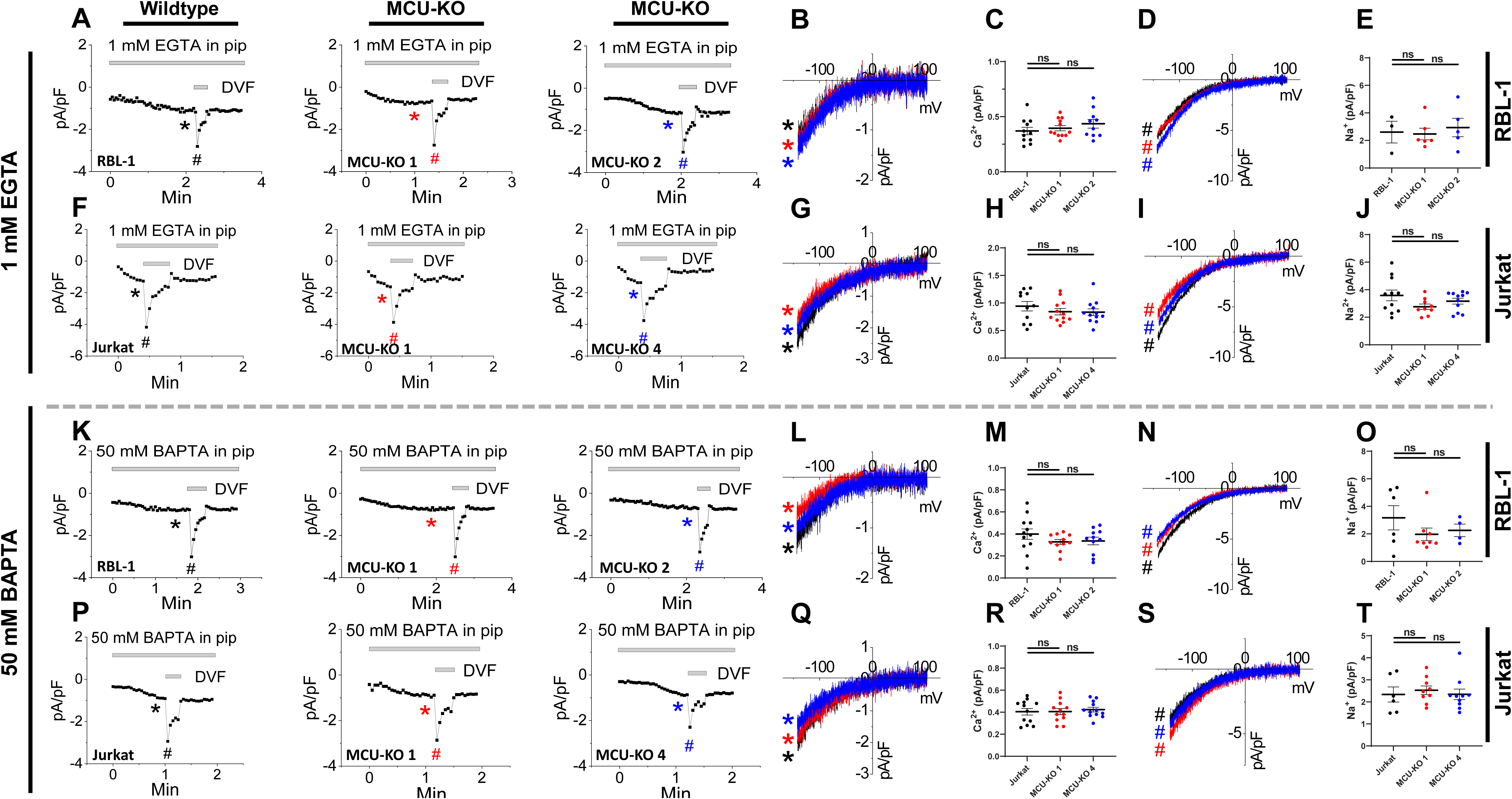
Effect of MCU-KO on CRAC currents. (**A**), CRAC current development taken at -100mV in (from left to right) parental RBL-1 mast cells, MCU-KO clone#1, and MCU-KO clone #2 of the same cells. Recordings were initiated after break-in with a pipette solution containing 1mM EGTA and a mitochondria-energizing cocktail (see methods). (**B, C**), I/V relationship taken from traces in (**A**) where indicated by color-coded asterisks (**B**) and quantification of peak CRAC current density (**C**) recorded in 20 mM Ca^2+^-containing bath solutions. (**D, E**), similar data to (**B, C**) but for Na^+^ CRAC currents recorded in divalent-free (DVF) solutions. (**F-J**), Similar recordings and data analysis to (**A-E**) for parental Jurkat T-cells and their MCU-KO clone #1 and clone #4. (**K-T**), Similar recordings and data analysis in parental and MCU-KO RBL-1 and Jurkat cells to (**A-J**) but using a pipette solution containing 50 mM BAPTA.

**Supplementary Figure 8.**
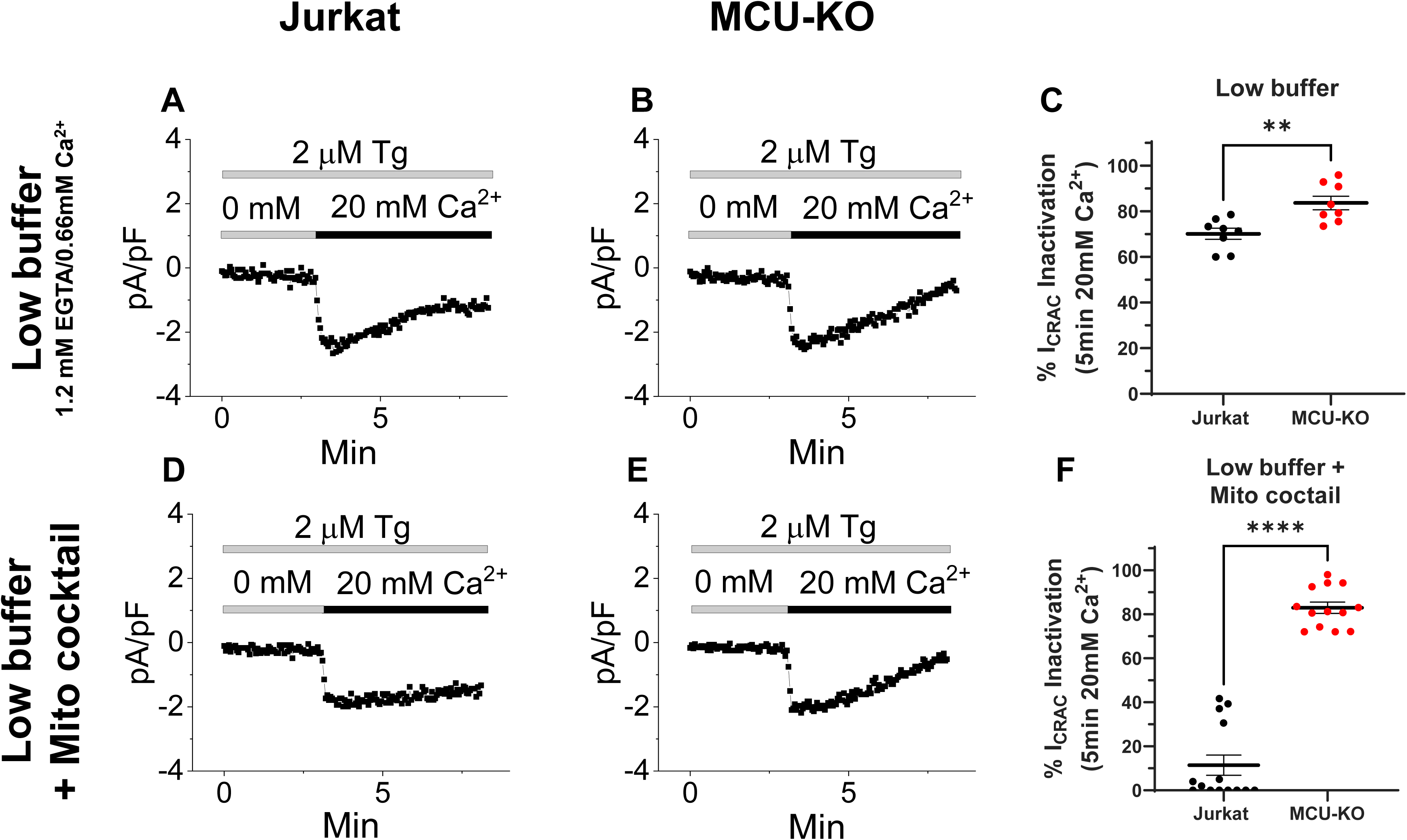
MCU-KO promotes slow CDI of CRAC currents. Slow CDI of CRAC currents recorded in parental Jurkat cells (**A, D**) and their MCU-KO counterparts (**B, E**) with a pipette solution containing 1.2 mM EGTA and 0.66 mM Ca^2+^ in either the presence (**D, E**) and absence (**A, B**) of the mitochondria-energizing cocktail. % of CDI is reported for all conditions in (**C, F**).

**Supplementary Figure 9.**
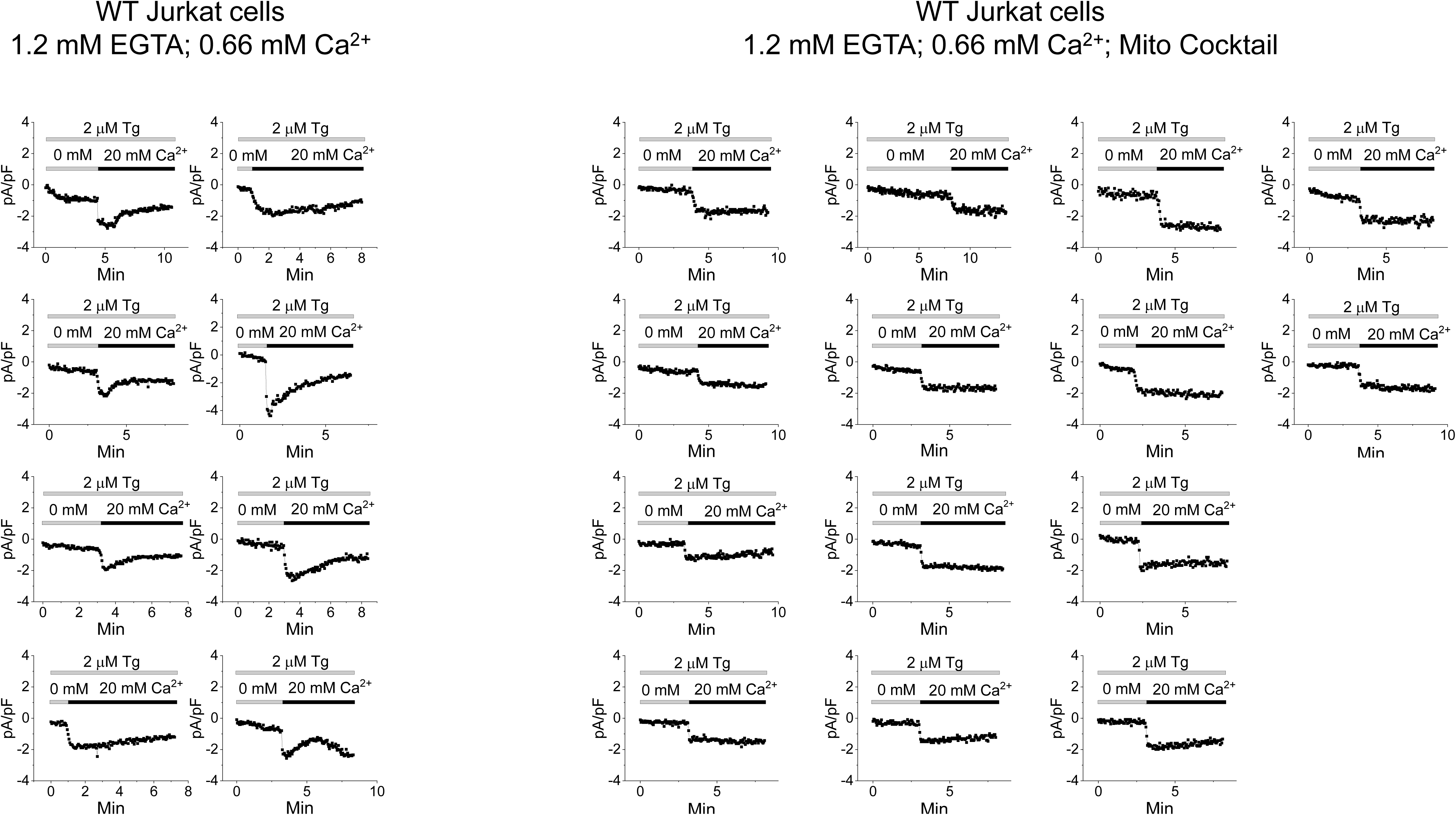
MCU-KO promotes slow CDI of CRAC currents. CDI recordings shown for all individual wildtype Jurkat cells.

**Supplementary Figure 10.**
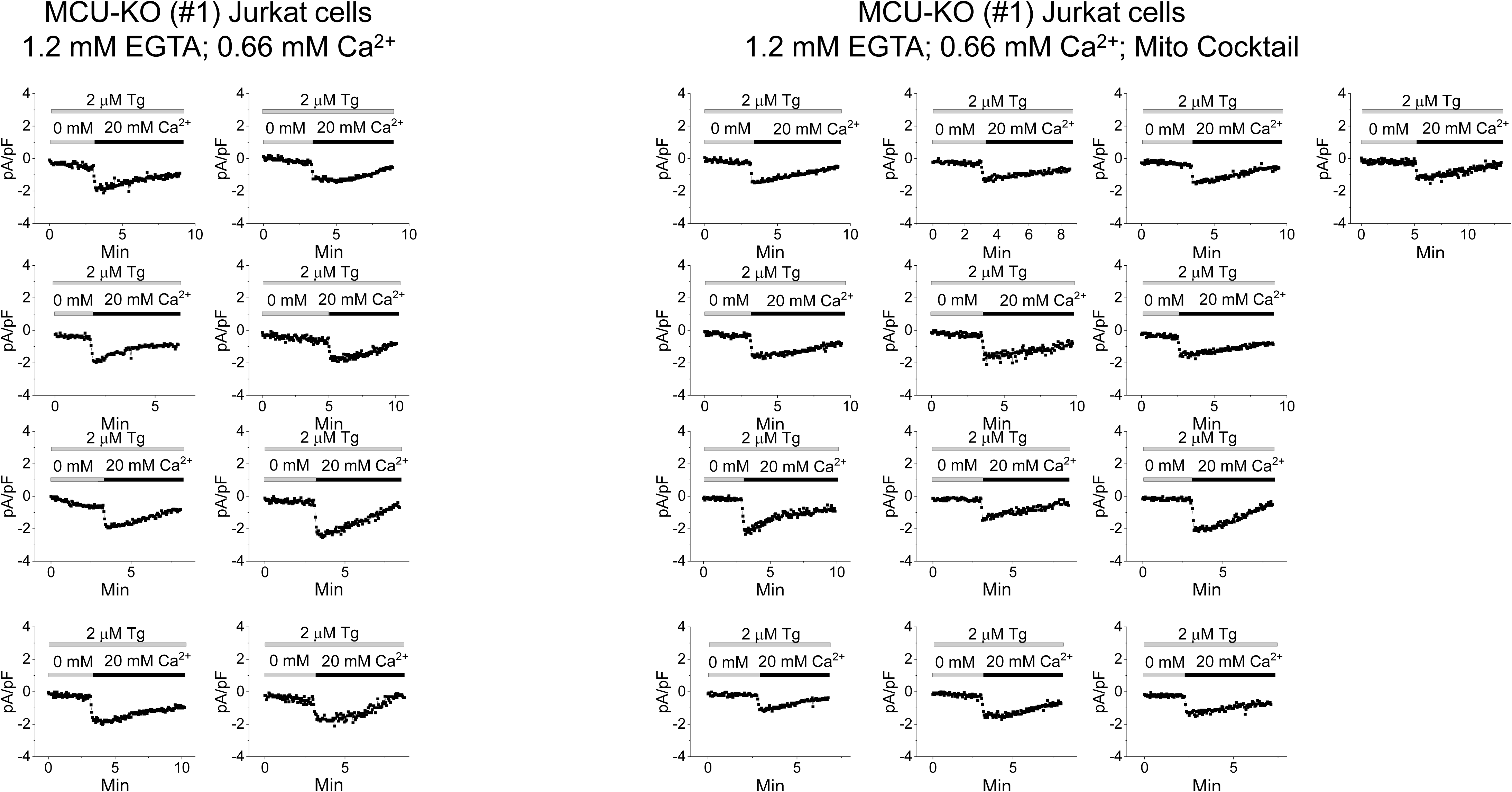
MCU-KO promotes slow CDI of CRAC currents. CDI recordings shown for all individual MCU-KO Jurkat cells.

**Supplementary Figure 11.**
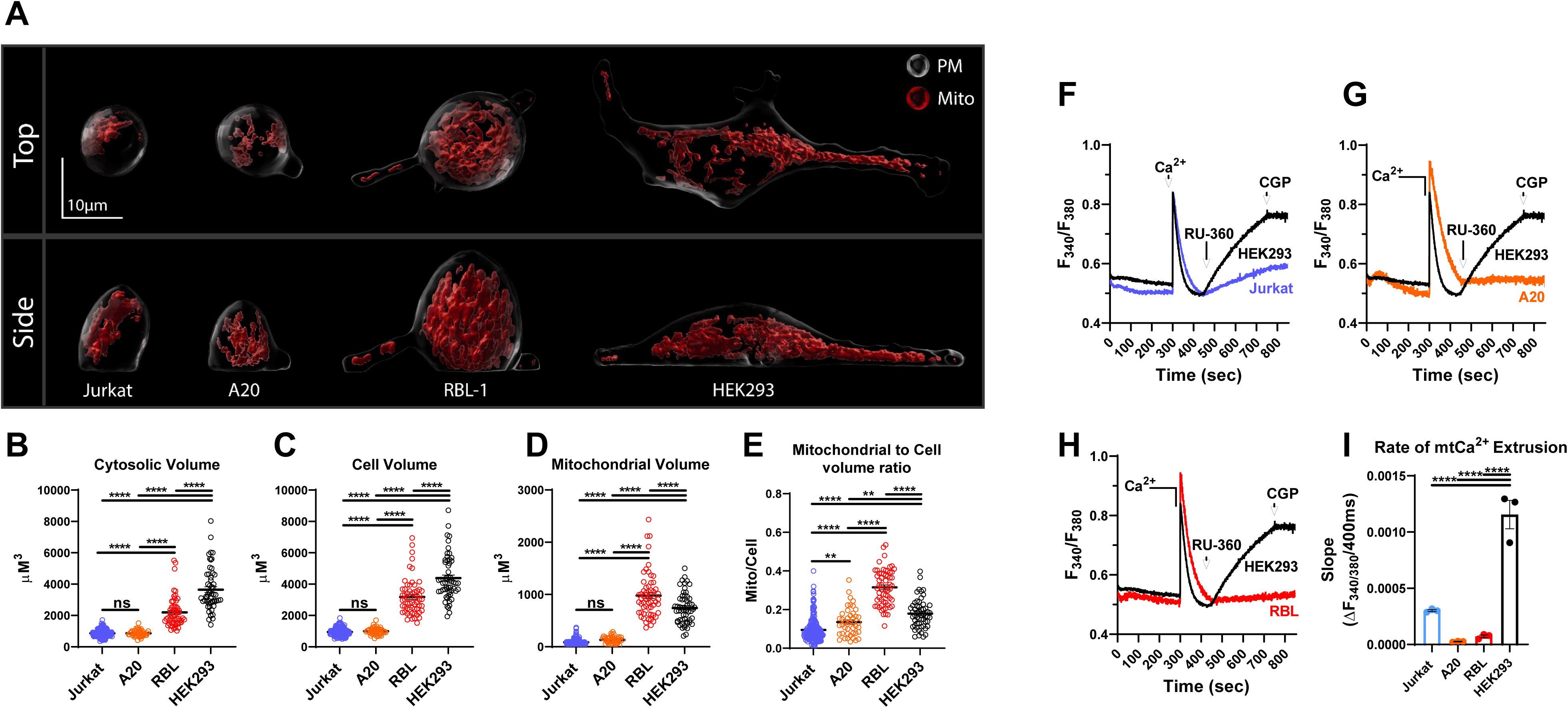
Cell and mitochondrial volume and Ca^2+^ extrusion differ between different cell lines. **(A)**, Representative three-dimensional rendering of mitochondrial volume (red) relative to total cell volume in wildtype Jurkat, A20, RBL-1 and HEK293 cells. Cytosolic (**B**), cell (**C**) and mitochondrial (**D**) volumes, and mitochondrial/cell volume ratio (**E**) were calculated and statistically analyzed. Mitochondrial Ca^2+^ extrusion was determined in HEK293 cells by comparison to Jurkat (**F**), A20 (**G**) and RBL-1 (**H**) cells in the permeabilized cell preparation. Bolus 10 µM Ca^2+^ was added to allow Ca^2+^ uptake, followed by addition of the MCU inhibitor RU360 (1 µM) to determine the rate of mitochondrial Ca^2+^ extrusion. The NCLX inhibitor, CGP37157 (10 µM) was added at the end of the recordings. Mitochondrial Ca^2+^ extrusion was calculated and statistically reported in (**I**).

**Supplementary Figure 12.**
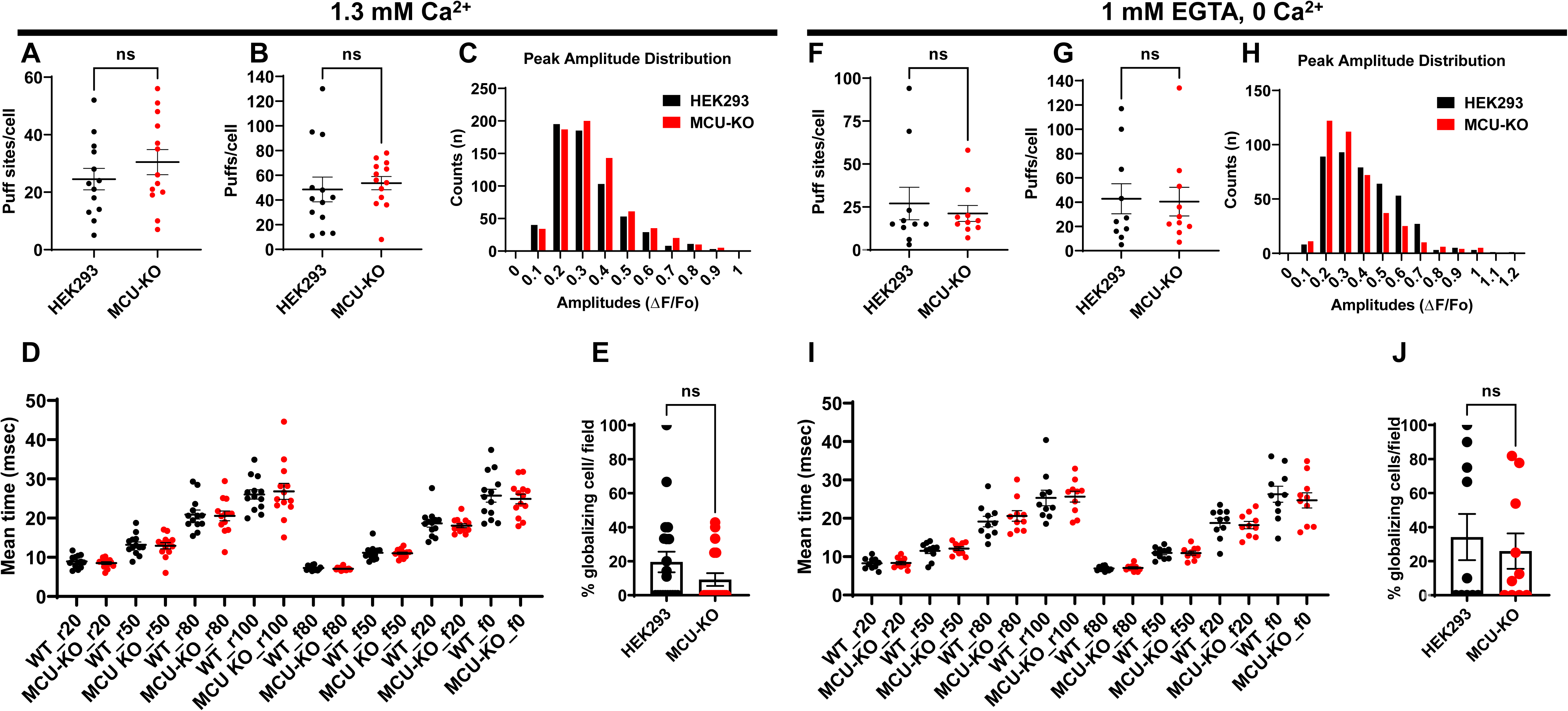
MCU-KO has no effect on IP_3_R-mediated Ca^2+^ puffs. (**A-E**) Ca^2+^ puffs were measured in Ca^2+^-containing buffer (1.3 mM Ca^2+^) using Cal-520 fluorescence ratios (ΔF/F_0_) from the center of single puff sites (1.3×1.3 µm) evoked by photolysis of ci-IP_3_ in wildtype HEK293 (n=20 independent experiments; 153 cells) and their MCU-KO counterparts (n=19 independent experiments; 132 cells). (**A, B**) 13 cells from each condition were randomly selected to quantify the number of puffs (**A**) and puff sites (**B**). (**C**) Amplitudes distribution of the Ca^2+^ puffs in wildtype HEK293 and MCU-KO cells. (**D**) Mean rise and decay times of fluorescence of Ca^2+^ puffs when it increases (r) or decreases (f) to 20%, 50%, 80%, and 100% from 13 cells each of HEK293 and MCU-KO. (**E**) Bar graph showing the proportions of wildtype HEK293 and MCU-KO cells in which the calcium signals globalized within 60 sec. (**F-J**), Similar experiments and data representation to (**A-E**) in wildtype HEK293 (n=11 independent experiments; 93 cells) and MCU-KO (n=12 independent experiments; 109 cells) where Ca^2+^ puffs were measured in Ca^2+^-free buffer supplemented with 1 mM EGTA.

**Supplementary Figure 13.**
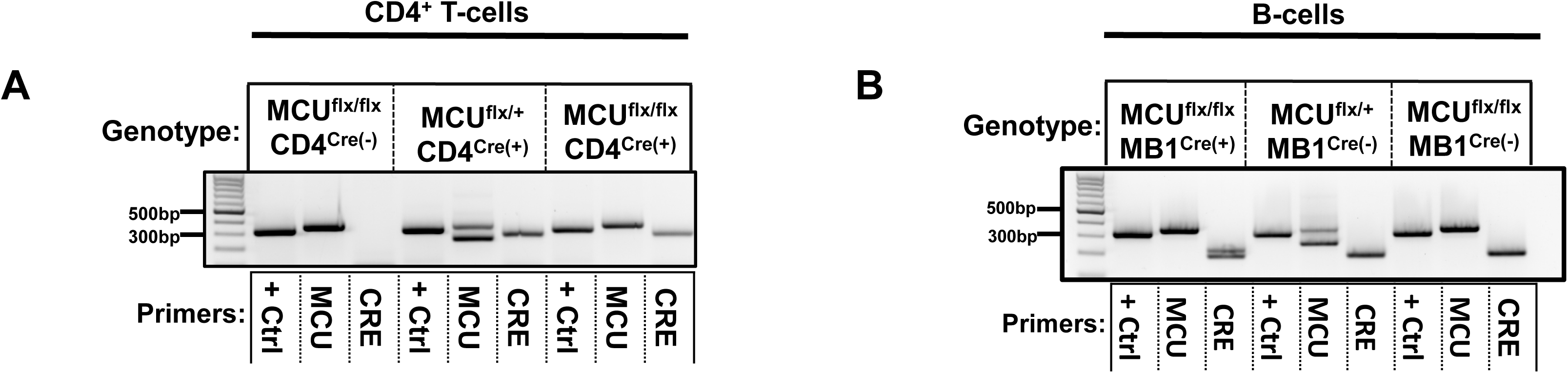
Generation of tissue-specific MCU-KD mice. MCU^flx/flx^ CD4^Cre(+)^ and MCU^flx/flx^ CD4^Cre(-)^ mice (**A**) and MCU^flx/flx^ MB1^Cre(+)^ and MCU^flx/flx^ MB1^Cre(-)^ mice (**B**) were identified by genotyping using specific primers as described in Methods.

**Supplementary Figure 14.**
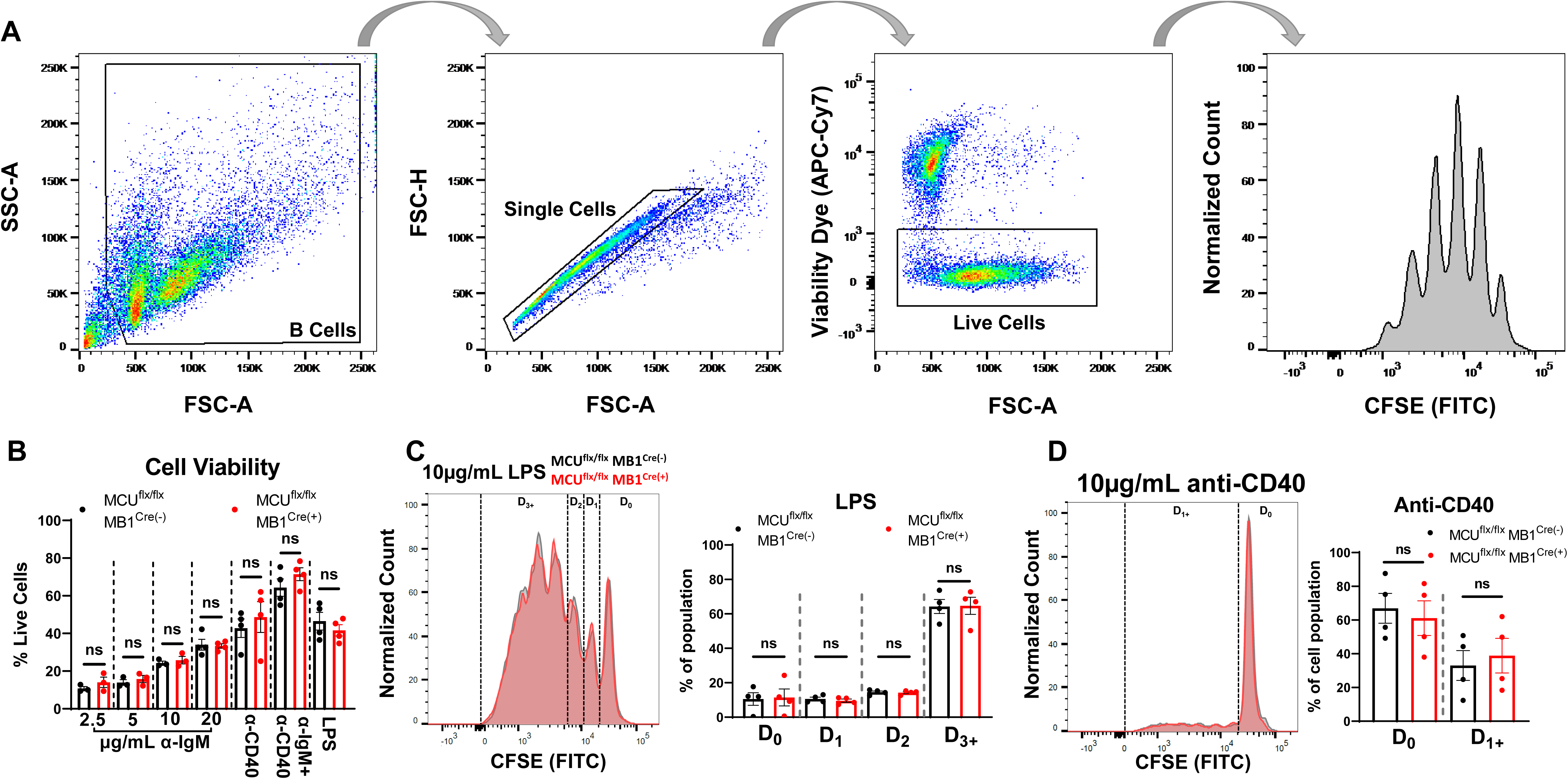
MCU-KD has no effect on LPS-mediated B-cell proliferation. **(A)**, Flow cytometry gating protocol used to assess B-cell proliferation. Intact and single B-cells are gated based on SSC and FSC signals. Of these cells, live cells are gated based on APC-cy7 staining and these live cells are used to quantify B-cell populations with different CFSE fluorescence. (**B**), B-cell viability is not different between MCU^flx/flx^ MB1^Cre(-)^ and MCU^flx/flx^ MB1^Cre(+)^ populations stimulated with anti-IgM, anti-IgM+anti-CD40, anti-CD40 or lipopolysaccharides (LPS). (**C**) B-cell proliferation in response to 10 µg/mL LPS determined by CFSE staining is not significantly different between MCU^flx/flx^ MB1^Cre(-)^ and MCU^flx/flx^ MB1^Cre(+)^ populations. (**D**), Stimulation with 10 µg/mL anti-CD40 alone did not induce significant B-cell proliferation in either MCU^flx/flx^ MB1^Cre(-)^ or MCU^flx/flx^ MB1^Cre(+)^ cells.

**Supplementary Figure 15.**
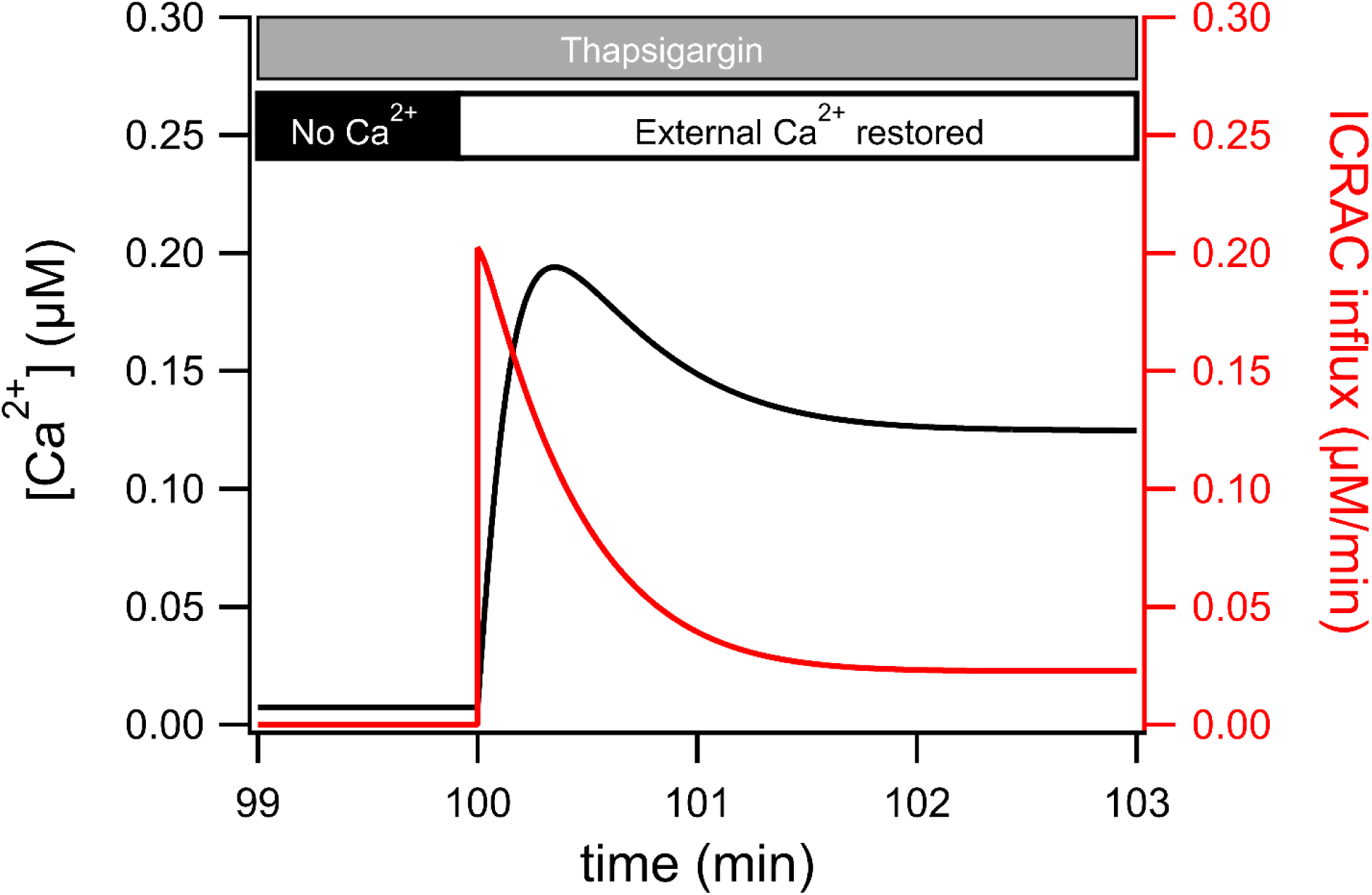
Model simulation of slow calcium-dependent inactivation of I_CRAC_. Ca^2+^ influx through I_CRAC_ (red curve plotted against the right-hand axis) and cytosolic Ca^2+^ concentration (black curve plotted against the left-hand axis) changes over time in a modeled cell treated first with thapsigargin in nominally free external Ca^2+^ until the ER was depleted. Upon restoration of external Ca^2+^ at 100 min, Ca^2+^ influx first increases quickly, then more slowly decays to a steady-state. The cytosolic Ca^2+^ concentration increases more slowly before reaching steady-state at a lower value. These results are qualitatively consistent with Fig. 2A of(*32*).

**Supplementary Figure 16.**
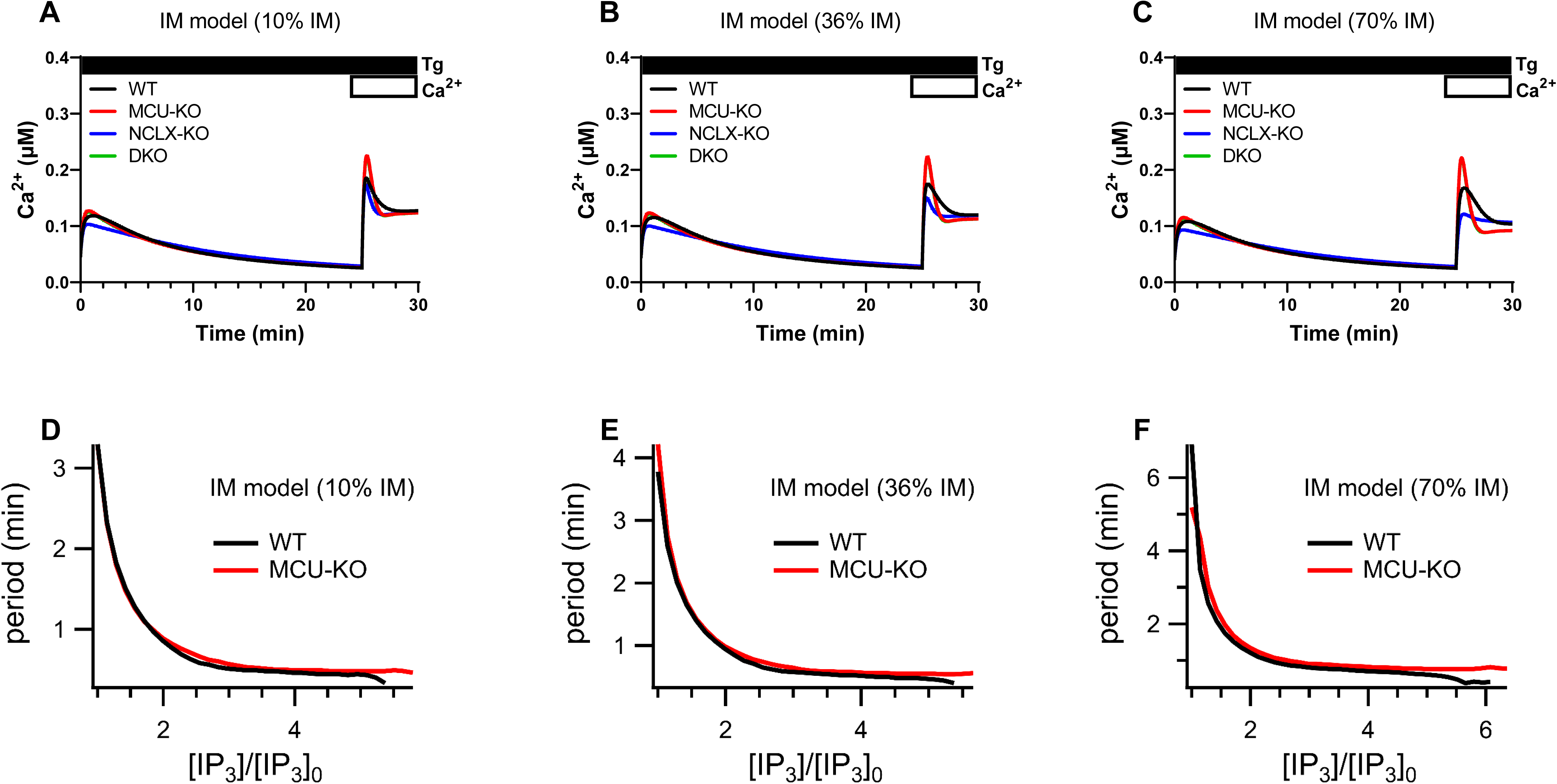
A Mathematical model of MCU-KO that also considers I_CRAC_ slow CDI. Models of I_CRAC_ microdomain (IM) between CRAC and MCU channels whereby slow CDI of CRAC channels is enhanced in the absence of MCU causing net reduction of I_CRAC_ across the PM. Variations of this model consider that either 10%, 36% or 70% of the incoming Ca^2+^ through I_CRAC_ enters mitochondria. (**A-C**). Model of the cytosolic Ca^2+^ signal in response to maximal store depletion with thapsigargin in the absence then presence of 2mM external Ca^2+^. (**E-G**), Period of Ca^2+^ oscillations (1/Frequency) as a function of relative concentrations of IP_3_ ([IP_3_]_0_ is the lowest value of [IP_3_] for which oscillations exist in the model with MCU) for the three different IM models from (**A-C**).

**Supplementary Table 1.**
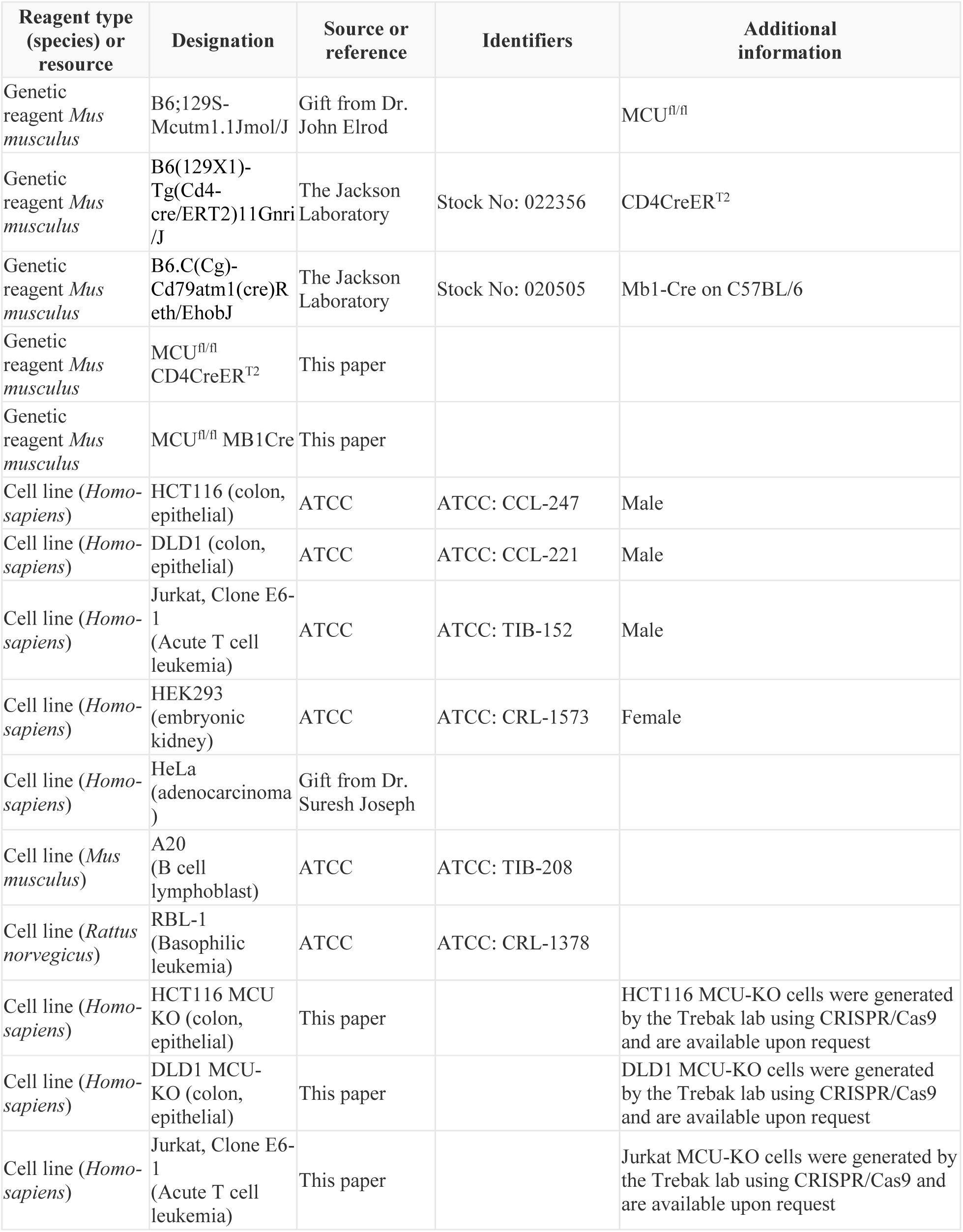
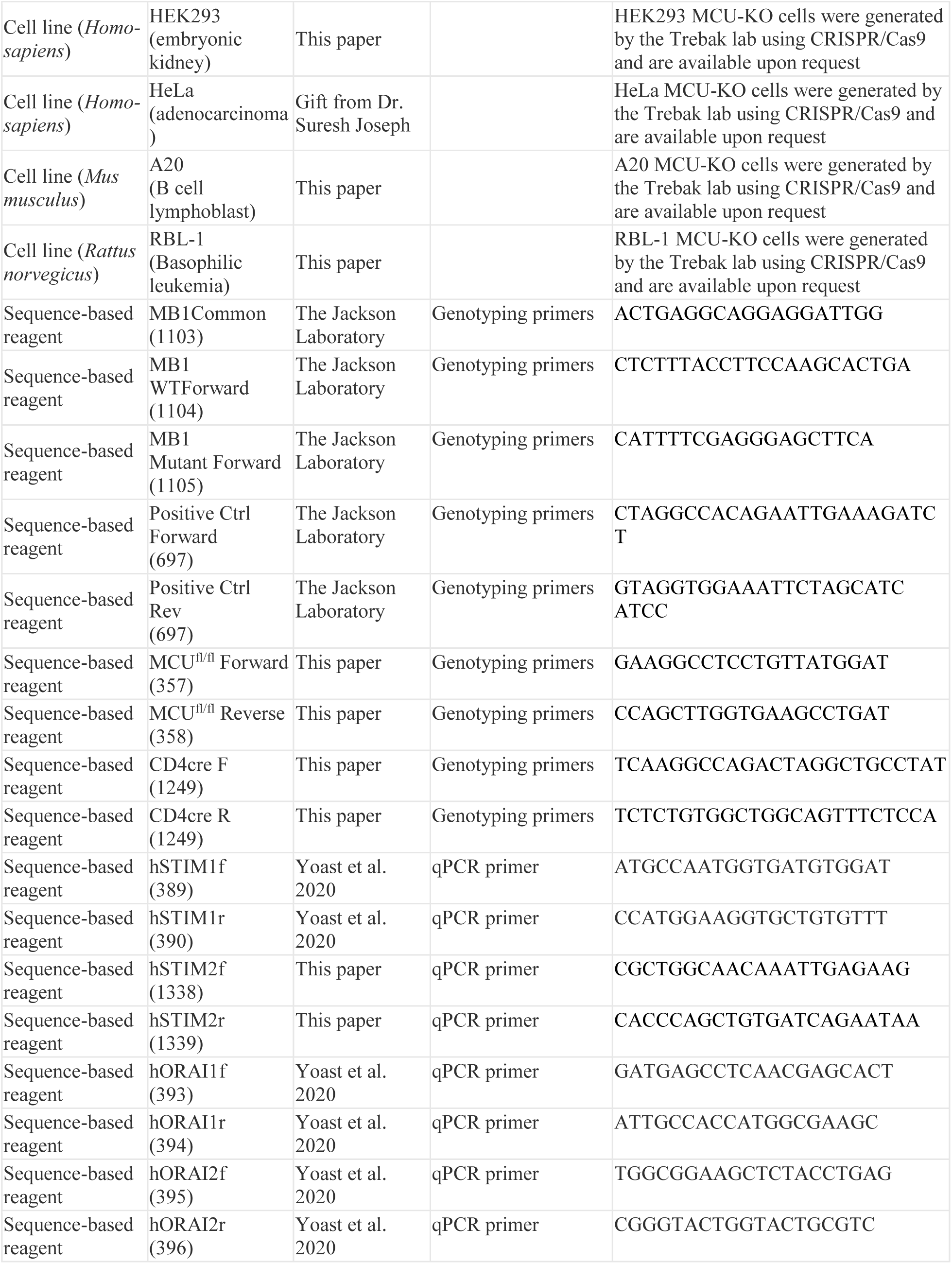
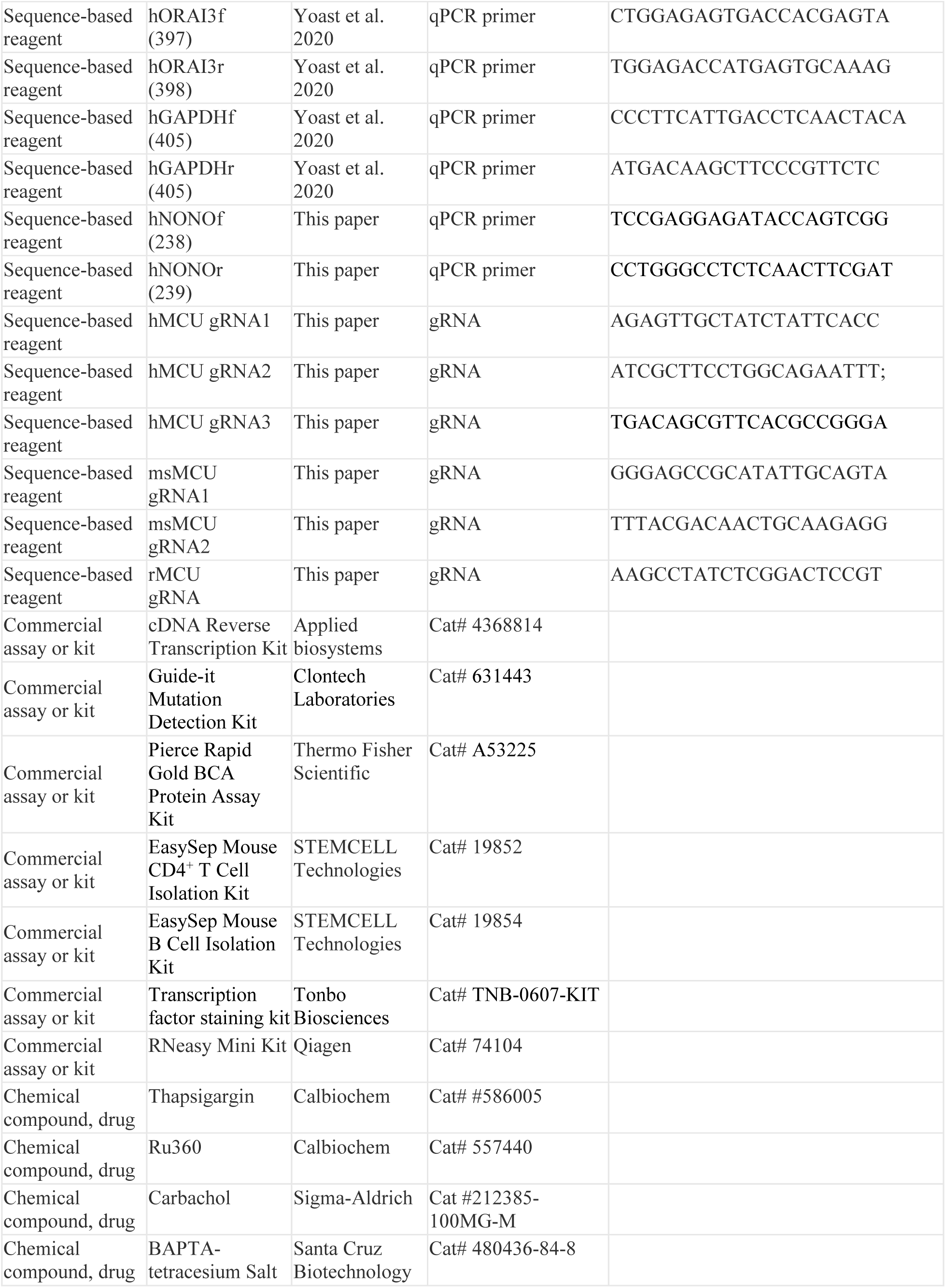
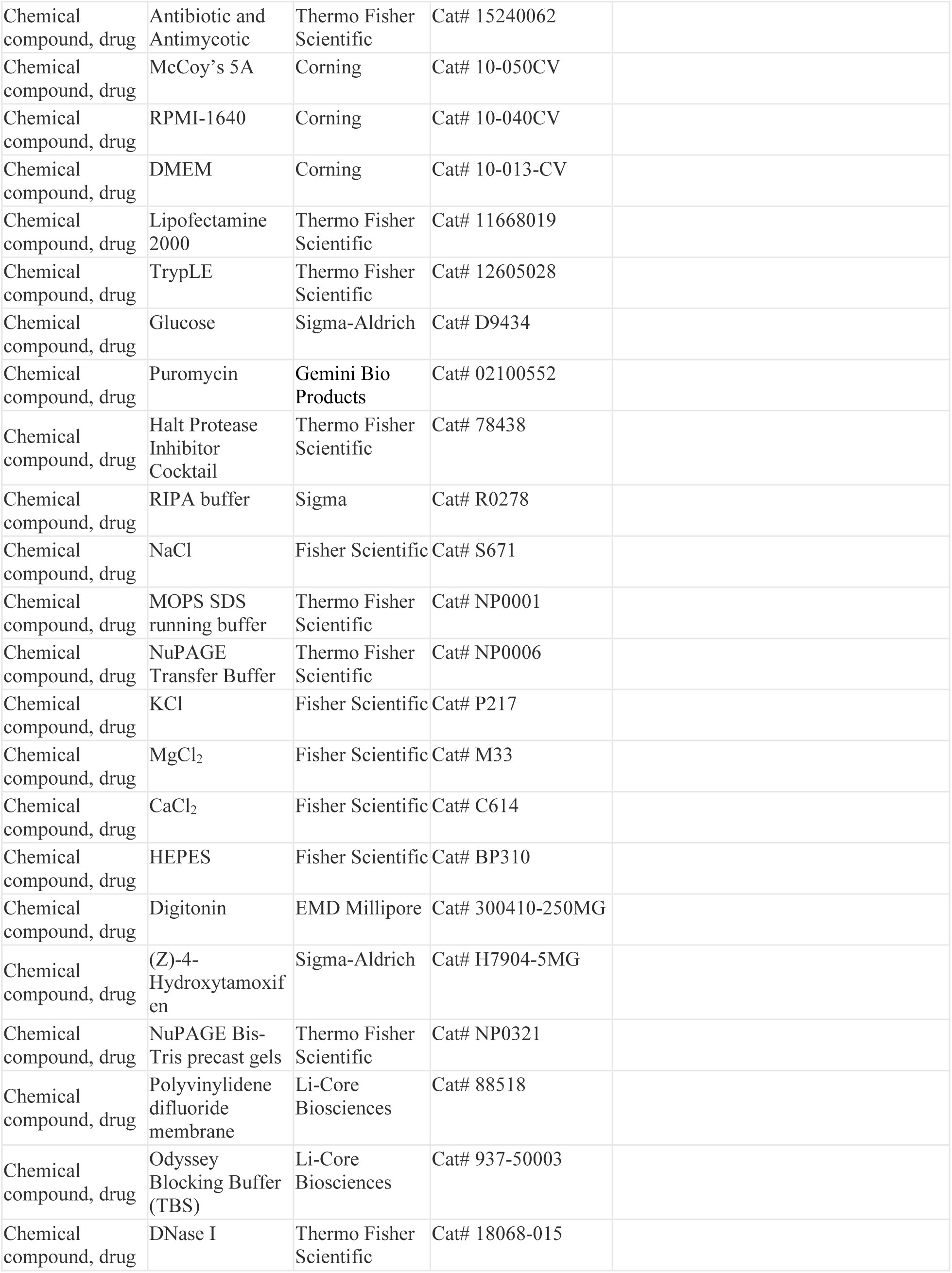
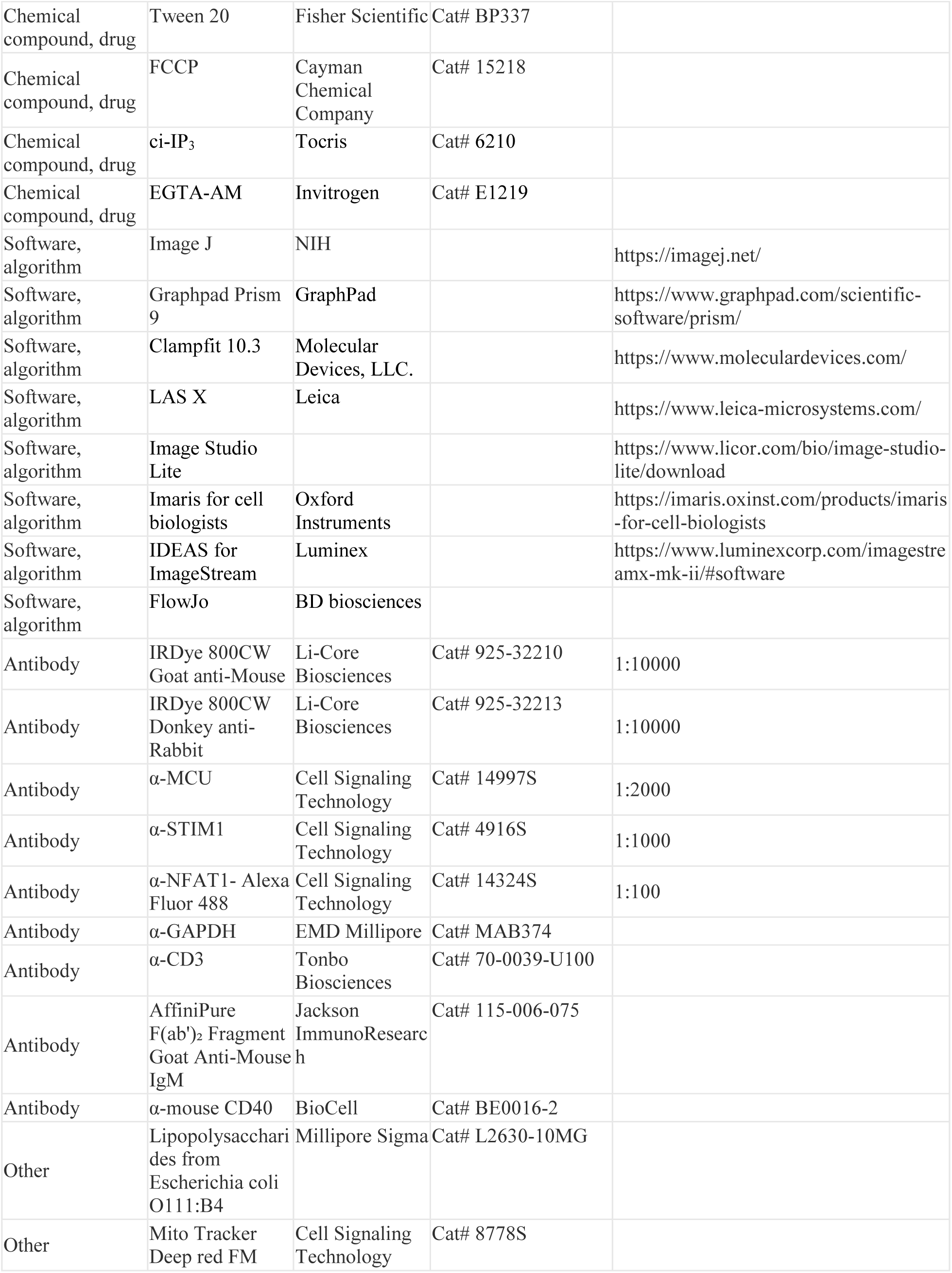
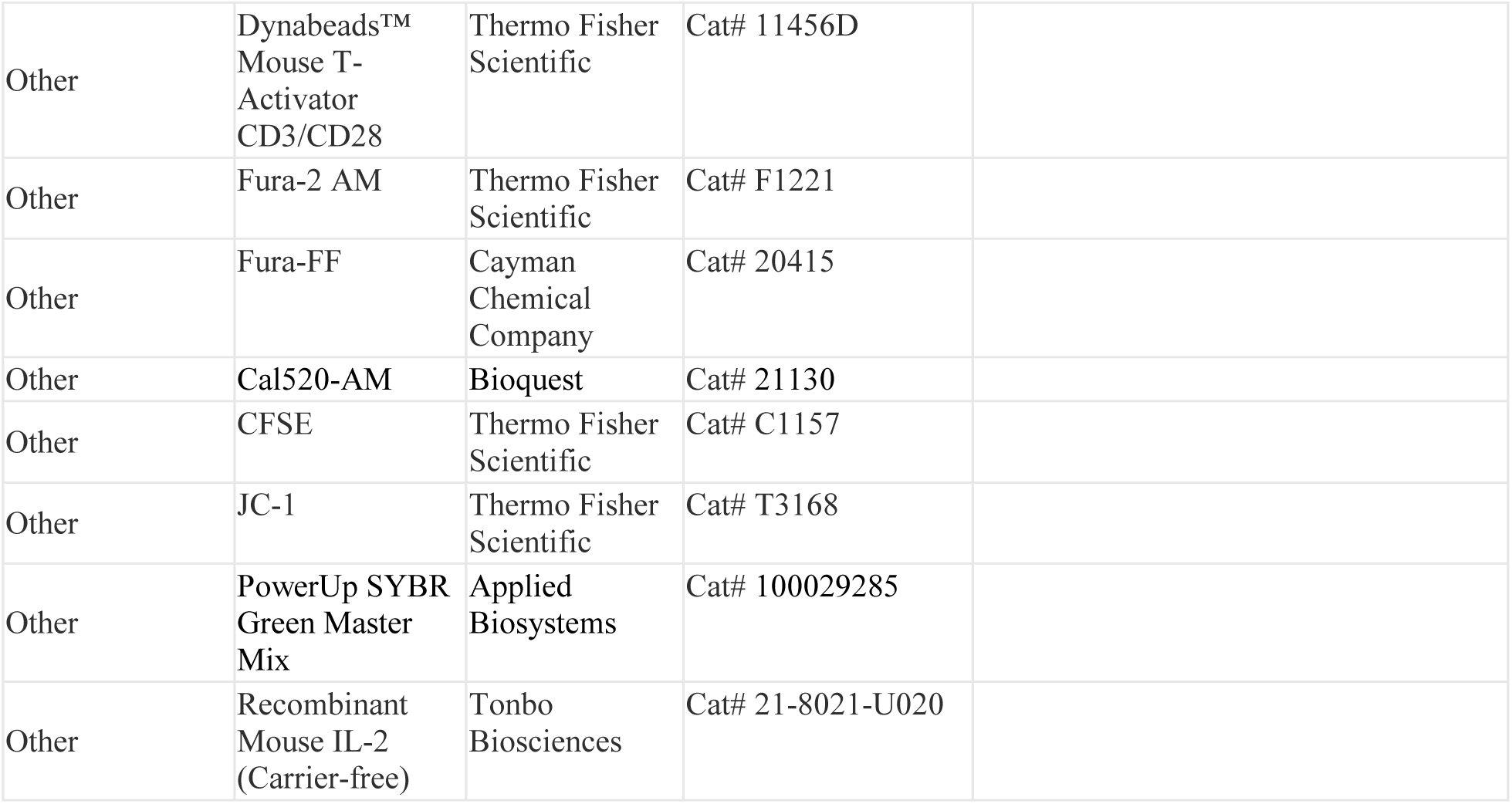
List and source of reagents, antibodies, chemicals, primers, gRNA, cell lines and mice used in the study.

